# Modeling phase separation dynamics of SynGAP and PSD-95 in postsynaptic densities

**DOI:** 10.64898/2026.07.23.740312

**Authors:** Sarah Sakly, Gregory D. Conradi Smith

## Abstract

SynGAP and PSD-95 undergo liquid–liquid phase separation in vitro, motivating models of the postsynaptic density (PSD) as a biomolecular condensate. However, how their association kinetics and intermolecular interactions shape activity-dependent partitioning remains unclear. We develop a multicomponent framework that couples equilibrium Flory–Huggins calculations in conserved total-composition space to four-species Cahn–Hilliard–reaction dynamics for free SynGAP, free PSD-95, the SynGAP–PSD-95 complex, and solvent. Complex formation and dissociation are represented by explicit forward and reverse activity-based mass-action rates. These calculations show that the equilibrium association constant and self- and cross-interaction energies jointly control the extent and composition of the two-phase region. For an illustrative LTP-like parameter switch, decreasing the equilibrium association constant and altering selected interaction energies redistributes SynGAP from the PSD-95-rich phase to the PSD-95-poor phase without eliminating the PSD-95-rich phase. A separate equilibrium comparison represents haploinsufficiency as a 50% reduction in total SynGAP. For the illustrative parameter sets, the reduced-SynGAP composition remains within the two-phase region but has different tie-line endpoints and phase compositions than the control composition.

## 1 Introduction

SynGAP is a highly abundant GTPase-activating protein (GAP) in the postsynaptic density (PSD) of excitatory synapses. By accelerating hydrolysis of GTP by Ras and Rap, Syn-GAP constrains signaling pathways downstream of NMDA receptors that promote synaptic strengthening and AMPA receptor insertion, thereby stabilizing basal synaptic efficacy [8, 11, 13].

NMDA receptor–mediated calcium influx induces kinase-dependent phosphorylation of SynGAP. Phosphorylation within regulatory domains modulates Ras-GAP activity, PSD localization, and AMPA receptor insertion [11, 24, 25]. Together, phosphorylation-dependent regulation and GAP activity allow SynGAP to link activity-dependent biochemical signaling to changes in synaptic strength during long-term potentiation (LTP).

Consistent with this central regulatory role, altered SynGAP expression has major effects on synaptic structure and function. Reduced SynGAP levels are associated with enlarged, prematurely stabilized dendritic spines, increased AMPA receptor content, and impaired capacity for subsequent LTP [13]. These synaptic abnormalities are particularly consequential during early postnatal life, when activity-dependent refinement of excitatory circuits is critical for cognitive and behavioral development. Reflecting this developmental sensitivity, pathogenic variants in *SYNGAP1* are predominantly loss-of-function, including protein-truncating mutations and mutations that disrupt protein stability, regulation, or localization [5, 11]. Disruption of SynGAP dosage or function is associated with a heterogeneous clinical spectrum including epilepsy, autism spectrum disorder, intellectual disability, developmental delay, and hypotonia [11, 13]. Understanding how SynGAP is retained within and redistributed from the PSD may therefore help connect changes in SynGAP dosage and regulation to altered synaptic function.

### 1.1 The postsynaptic density as a biomolecular condensate

The PSD serves as a central hub for synaptic signaling and plasticity, enabling rapid and spatially constrained biochemical interactions [22]. The scaffolding protein PSD-95 (SAP90) is a core organizational element of the PSD. Its multiple protein–protein interaction domains promote the assembly of signaling complexes, including through direct binding to SynGAP [16]. *In vitro* experiments have shown that multivalent interactions between PSD-95 and SynGAP can drive liquid–liquid phase separation (LLPS) [27]. These findings motivate condensate models of the PSD, although they do not by themselves establish condensate behavior of the intact PSD in vivo. Such models have been proposed to explain how the PSD remains spatially confined while preserving molecular mobility and the capacity for rapid reorganization in response to synaptic activity [17].

### 1.2 Molecular basis of SynGAP–PSD-95 interactions

PSD-95 contains three PDZ domains that mediate interactions with a broad array of synaptic proteins [14]. SynGAP localization and regulation within the PSD depend critically on binding between its C-terminal PDZ-binding motif and the PDZ domains of PSD-95 [11]. SynGAP–PSD-95 binding is dynamically regulated by phosphorylation. Kinases such as CaMKII and PLK2 reduce SynGAP affinity for PSD-95, promoting SynGAP displacement from the PSD and enabling competing PDZ-binding proteins, including AMPA receptor– associated proteins, to engage PSD-95 [24]. Experiments further showed that SynGAP directly competes with AMPA receptor–TARP complexes for condensate formation with PSD-95 and that this competition does not require SynGAP catalytic activity [2]. Calcium/calmodulin binding provides an additional regulatory layer by selectively weakening the interaction of SynGAP with the PDZ3 domain of PSD-95 [24]. Beyond regulating pairwise binding, these phosphorylation- and calcium-dependent mechanisms may influence the emergent material properties of the PSD.

### 1.3 Flory–Huggins modeling of biomolecular condensates

Flory–Huggins theory provides a mean-field description of mixing in polymer–solvent and multicomponent systems [10, 12]. For a binary polymer–solvent mixture, the bulk free-energy density is

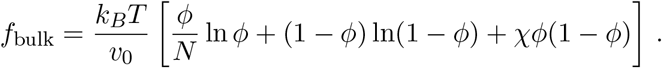

Here, *N* is the number of lattice sites of volume *v*_0_ occupied by one polymer molecule, *ϕ* is the polymer volume fraction, and 1 − *ϕ* is the solvent volume fraction. The logarithmic terms describe the entropy of mixing, whereas the dimensionless Flory–Huggins interaction parameter χ describes energetic deviations from ideal mixing. For sufficiently unfavorable interactions, the free-energy density can support coexistence between polymer-poor and polymer-rich phases, denoted *α* and *β*, with polymer volume fractions *ϕ*^*α*^ and *ϕ*^*β*^, respectively [20].

To introduce multicomponent phase-separation models for biomolecular condensates, consider a ternary Flory–Huggins mixture consisting of SynGAP (*S*), PSD-95 (*P*), and solvent (*Q*). The volume fractions satisfy *ϕ*_*s*_ + *ϕ*_*p*_ + *ϕ*_*q*_ = 1, and nonideal mixing is described by interaction parameters χ_*s,p*_, χ_*s,q*_, and χ_*p,q*_. In this case, the entropic and enthalpic contributions to the bulk free energy are

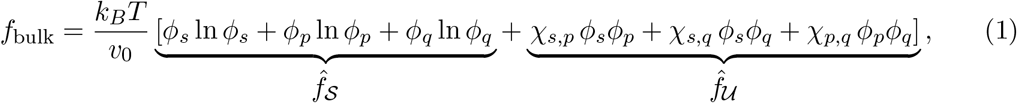

where *ϕ*_*q*_ = 1 − *ϕ*_*s*_ − *ϕ*_*p*_ and *N*_*s*_ = *N*_*p*_ = *N*_*q*_ = 1 for simplicity. We analyze this system using ternary phase diagrams, which show *f*_bulk_ over the (*ϕ*_*s*_, *ϕ*_*p*_, *ϕ*_*q*_) composition simplex (Fig. 1).

**Figure 1.**
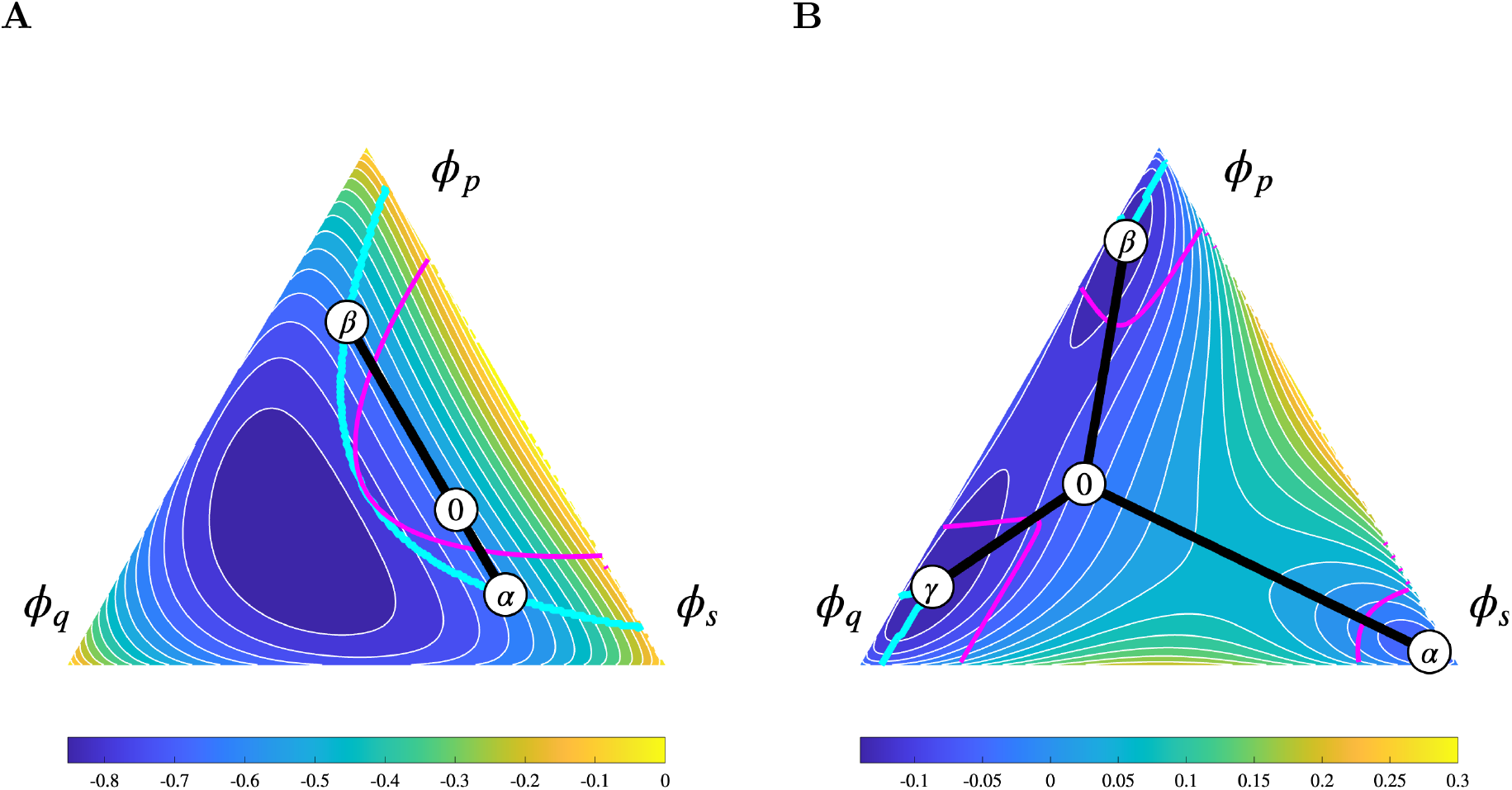
Ternary Flory–Huggins phase diagrams for the bulk free-energy density in Eq. 1, evaluated on a 200-point-per-axis simplex mesh. **A**: For χ_*s,p*_ = 3, χ_*s,q*_ = χ_*p,q*_ = 0, the overall composition ***ϕ***^0^ = (0.5, 0.3, 0.2) separates into phases with compositions ***ϕ***^*α*^ and ***ϕ***^*β*^. **B**: For χ_*s,p*_ = 3.95, χ_*s,q*_ = 3.62, and χ_*p,q*_ = 2.56, the overall composition ***ϕ***^0^ = (0.2, 0.35, 0.45) separates into phases with compositions ***ϕ***^*α*^, ***ϕ***^*β*^, and ***ϕ***^*ϕ*^. In both panels, the white circle labeled 0 indicates ***ϕ***^0^. The labeled coexistence compositions are obtained from the lower-convex-hull facet containing ***ϕ***^0^; the black segments connect them, and the corresponding phase fractions reconstruct ***ϕ***^0^ by the lever rule (Eq. 34). Cyan and magenta curves indicate the binodal and spinodal boundaries, respectively.

These examples illustrate twoand three-phase coexistence in a ternary mixture. The binodal bounds the region in which phase coexistence lowers the free energy relative to the homogeneous state, whereas the spinodal marks the loss of local stability of the homogeneous state. In the metastable region between the binodal and spinodal, phase separation requires nucleation by a finite-amplitude fluctuation [6, 20]. Each tie line joins two coexisting equilibrium compositions, so its orientation indicates the direction of compositional partitioning between the phases. With *ϕ*_*q*_ at the lower left, *ϕ*_*s*_ at the lower right, and *ϕ*_*p*_ at the top, a nearly horizontal tie line primarily reflects exchange between solvent and SynGAP with little change in PSD-95, whereas an upward-directed tie line indicates stronger partitioning of PSD-95. Parallel tie lines indicate a similar partitioning direction across the coexistence region, while rotating or fanning tie lines indicate composition-dependent partitioning. Tie-line length measures the compositional contrast between phases, and their relative amounts follow from the lever rule.

### 1.4 Cahn–Hilliard modeling of phase separation

Figs. 2 and 3 show Cahn–Hilliard simulations of the (*ϕ*_*s*_, *ϕ*_*p*_, *ϕ*_*q*_) ternary system for the two- and three-phase cases in Fig. 1A and B, respectively. Whereas the Flory–Huggins free energy determines the equilibrium phase compositions, the Cahn–Hilliard equations describe the conserved spatial redistribution of the components toward those states. We eliminate the solvent field using conservation of volume, *ϕ*_*q*_ = 1 − *ϕ*_*s*_ − *ϕ*_*p*_, and define the solvent-referenced chemical-potential differences

**Figure 2.**
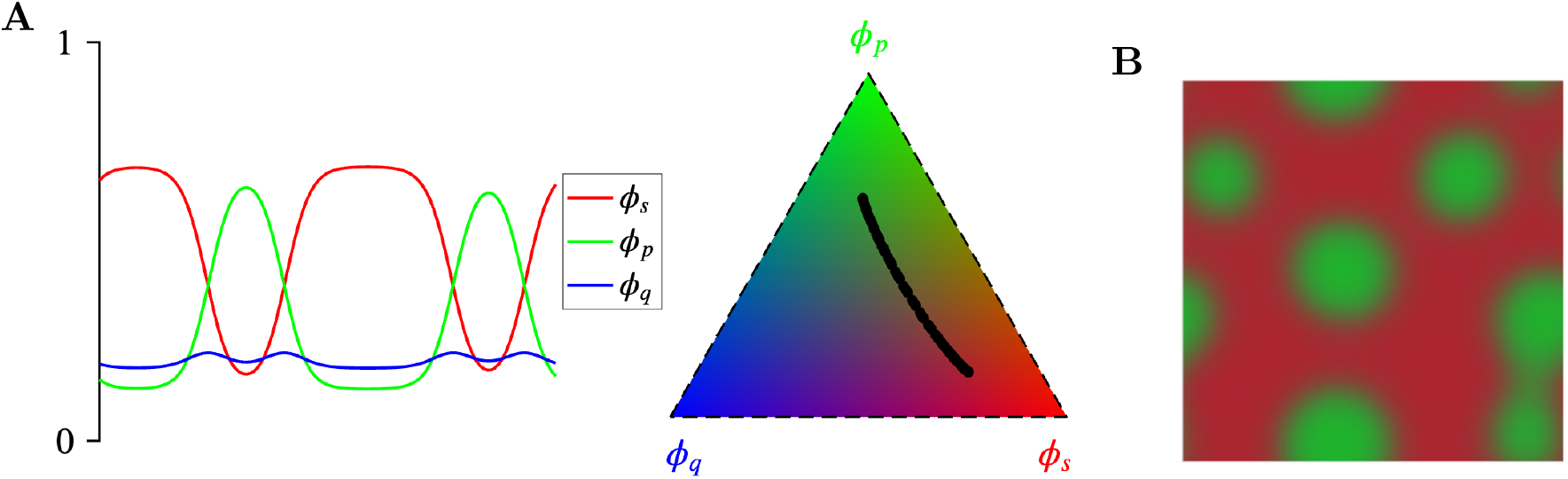
Two-phase Cahn–Hilliard simulations. **A**: One-dimensional simulation. **B**: Two-dimensional simulation. Spatially homogeneous initial states with random perturbations (not shown) undergo spinodal decomposition into two phases, with the coexistence compositions labeled *β* and *γ* in the ternary diagram of Fig. 1A. A two-field semi-implicit spectral method evolves (*ϕ*_*s*_, *ϕ*_*p*_), with *ϕ*_*q*_ = 1 − *ϕ*_*s*_ − *ϕ*_*p*_. Each simulation uses 128 grid points per spatial direction and a domain length of 40, #*x* = 0.3125, #*t* = 3 × 10^−4^, and final time *t* = 300. The Flory–Huggins interaction parameters χ_*ij*_ are as in Fig. 1A.

**Figure 3.**
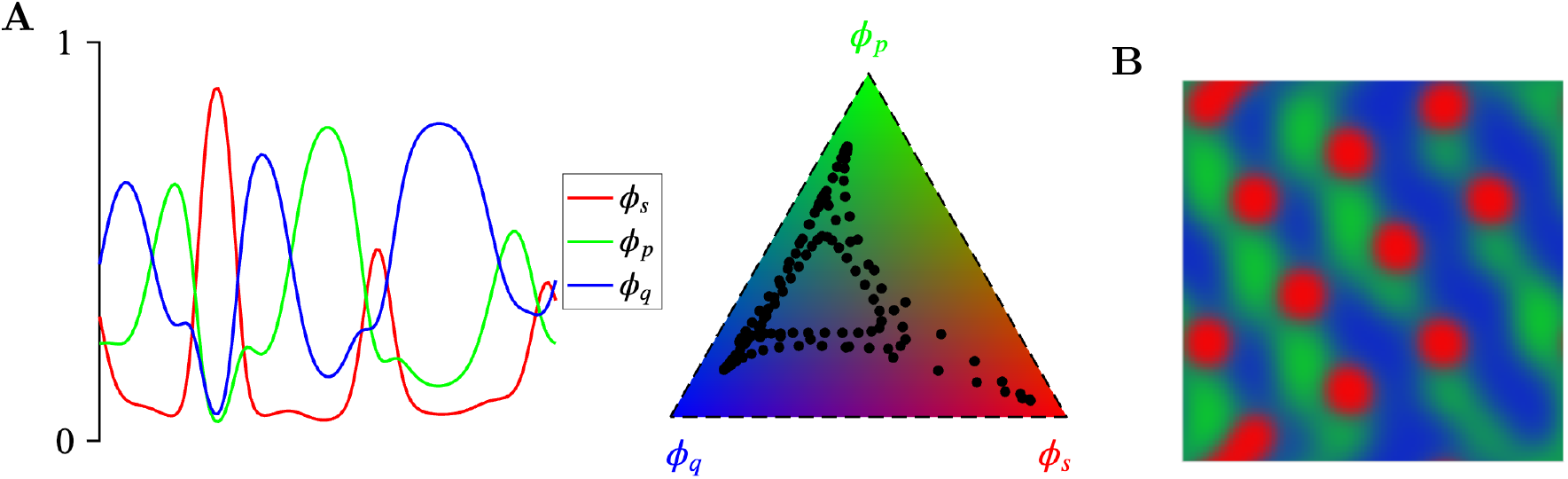
Three-phase Cahn–Hilliard relaxation. **A**: One-dimensional simulation. **B**: Two-dimensional simulation. The simulations begin from deterministic, smoothed multidomain fields constructed from the grid-resolved coexistence compositions labeled *α, γ*, and *ϕ* in Fig. 1B. Both panels show snapshots at *t* = 6. The domain length, Δ*x*, and Δ*t* are as in Fig. 2; the interaction parameters χ_*ij*_ are as in Fig. 1B.

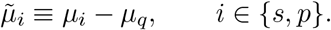

Each chemical-potential difference comprises bulk and gradient contributions,

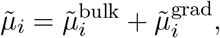

where

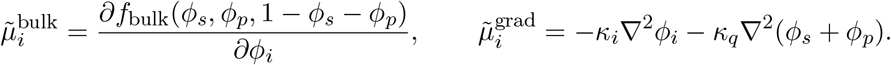

The resulting two-field Cahn–Hilliard system is

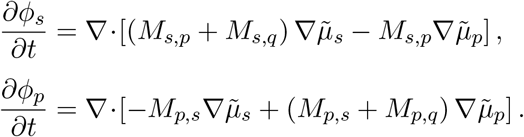

For these preliminary simulations, we set *κ*_*i*_ = 1 for *i* ∈ *{s, p, q}* and *M*_*i,j*_ = 1*/*3 for *i ≠ j*.

Fig. 2A shows a snapshot at *t* = 300 of one-dimensional spatial profiles after spinodal decomposition into PSD-95-rich and PSD-95-poor phases (green curve). The Cahn–Hilliard fields approach the lower-hull estimates

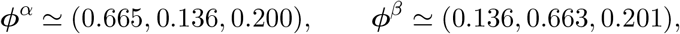

consistent with the equilibrium predicted from the Flory–Huggins free-energy density at overall composition ***ϕ***^0^ (Fig. 1A). Mapping the spatial profiles ***ϕ***(*x, t*) (filled black circles) onto the ternary plot shows their approach to these coexistence compositions (Fig. 2A, triangle). Fig. 2B repeats the Cahn–Hilliard calculation in two spatial dimensions with periodic boundary conditions.

Fig. 3 applies the same two-field Cahn–Hilliard model to the bulk free energy in Fig. 1B, with ***ϕ***^0^ = (0.20, 0.35, 0.45) and (χ_*s,p*_, χ_*s,q*_, χ_*p,q*_) = (3.95, 3.62, 2.56). Because the local instability is weak at this mean composition, we initialize smoothed regions near three lower-hull compositions: ***ϕ***^*α*^ ≃ (0.940, 0.025, 0.035), ***ϕ***^*β*^ ≃ (0.035, 0.819, 0.146), and ***ϕ***^*ϕ*^ ≃ (0.045, 0.151, 0.804). The one- and two-dimensional profiles at *t* = 6 retain multiple regions of each phase while their interfaces relax (Fig. 3A,B), consistent with the three-phase coexistence shown in Fig. 1B.

### 1.5 Organization of the paper

The remainder of the main text is organized as follows. (i) We formulate a four-species model that relates total composition to equilibrium species fractions, constructs the Flory–Huggins bulk free energy and activity-based reaction kinetics, and derives Cahn–Hilliard–reaction equations for spatial evolution in an idealized domain (§2). (ii) We examine how equilibrium association strength, relative molecular volumes, and heterotypic and homotypic interaction energies, formulated following Lin *et al*. [17], shape equilibrium phase behavior and protein partitioning between PSD-95-poor and PSD-95-rich phases (§3.1 and 3.2). (iii) Using a semi-implicit spectral method, we numerically integrate the Cahn–Hilliard–reaction equations to model SynGAP dispersal in response to an LTP-like switch in the equilibrium association constant *K* and selected interaction energies (§3.3 and 3.4). (iv) We use equilibrium phase diagrams to examine how a 50% reduction in conserved total SynGAP changes pre- and post-switch phase coexistence (§3.5). (v) We conclude by discussing model limitations, implications for SynGAP haploinsufficiency, and directions for future work (§4).

## 2 Model formulation

We model SynGAP–PSD-95 LLPS as a multicomponent Cahn–Hilliard–reaction system comprising four species volume-fraction fields. Association and dissociation are modeled using activity-based mass-action rates that incorporate nonideal bulk Flory–Huggins interactions. The free-energy functional includes bulk and gradient contributions [7],

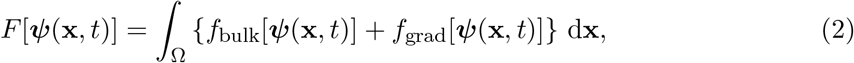

where Ω is the periodic one- or two-dimensional simulation domain and ***ψ*** = (*ψ*_*s*_, *ψ*_*p*_, *ψ*_*sp*_, *ψ*_*q*_) denotes the vector of volume fractions of free SynGAP, free PSD-95, the SynGAP–PSD-95 complex, and solvent, respectively. We impose local incompressibility, so the species volume fractions satisfy

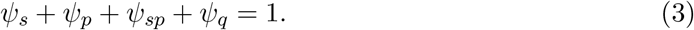

We model the reversible association of SynGAP and PSD-95 as

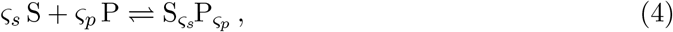

where S denotes free SynGAP, P denotes free PSD-95, 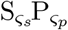 denotes the SynGAP–PSD-95 complex, and *ς*_*s*_ and *ς*_*p*_ are stoichiometric coefficients specifying the numbers of SynGAP and PSD-95 molecules in each complex, respectively. The reaction redistributes protein between free and bound species but conserves total SynGAP and total PSD-95. We express the corresponding conserved protein volume fractions as *ϕ*_*s*_ = *ψ*_*s*_ + *σ*_*s*_*ψ*_*sp*_, *ϕ*_*p*_ = *ψ*_*p*_ + *σ*_*p*_*ψ*_*sp*_, with *σ*_*s*_ + *σ*_*p*_ = 1. The coefficients *σ*_*s*_ = *ς*_*s*_*N*_*s*_*/N*_*sp*_ and *σ*_*p*_ = *ς*_*p*_*N*_*p*_*/N*_*sp*_ are the fractions of the complex volume attributable to SynGAP and PSD-95, respectively. Here, *N*_*s*_ and *N*_*p*_ are the relative molecular volumes of free SynGAP and PSD-95, and *N*_*sp*_ = *ς*_*s*_*N*_*s*_ + *ς*_*p*_*N*_*p*_ is the corresponding relative molecular volume of the SynGAP–PSD-95 complex.

### 2.1 Equilibrium species fractions

In equilibrium Flory–Huggins calculations, the species and total volume fractions are related by

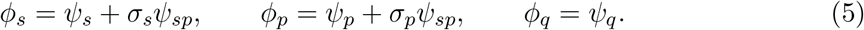

The following schematic summarizes this map:

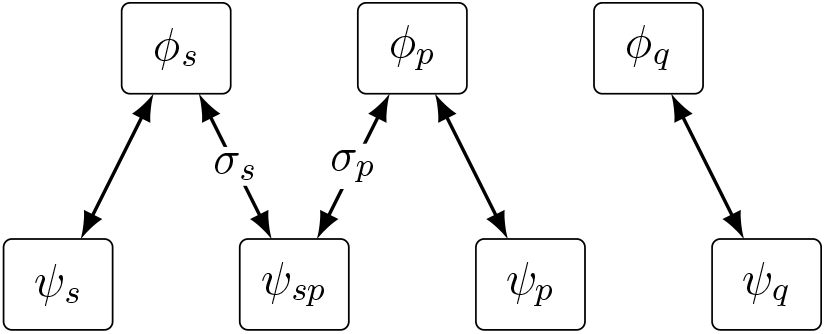

The species coordinates satisfy Eq. 3, whereas the total-composition coordinates satisfy *ϕ*_*s*_ + *ϕ*_*p*_ + *ϕ*_*q*_ = 1. Although *ψ*_*q*_ = *ϕ*_*q*_, we retain both symbols to distinguish the species and total-composition coordinate systems. For each total composition ***ϕ*** = (*ϕ*_*s*_, *ϕ*_*p*_, *ϕ*_*q*_), we determine the equilibrium species fractions by minimizing the bulk free energy subject to Eq. 5 and the nonnegativity of all species fractions. Substituting these fractions into *f*_bulk_ yields the reduced ternary Flory–Huggins free-energy surface used to construct the phase diagrams.

### 2.2 Flory–Huggins bulk free energy density

Following the equilibrium model of Lin *et al*. [17], we decompose the bulk free-energy density into entropic, enthalpic, and reaction contributions

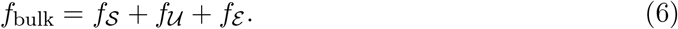

#### Entropy (*S*)

The term *f*_*S*_ in Eq. 6 represents the entropy of mixing among free SynGAP (*ψ*_*s*_), free PSD-95 (*ψ*_*p*_), the SynGAP–PSD-95 complex (*ψ*_*sp*_), and solvent (*ψ*_*q*_). Using the standard Flory–Huggins lattice expression, we write

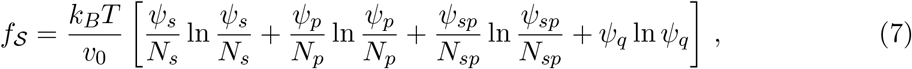

where *N*_*i*_ = *v*_*i*_*/v*_0_ is the relative molecular volume of species *i* in units of the lattice-site volume *v*_0_.

#### Enthalpy (*U*)

The Flory–Huggins enthalpy *f*_*U*_ accounts for pairwise interactions among free SynGAP, free PSD-95, the SynGAP–PSD-95 complex, and solvent and is given by

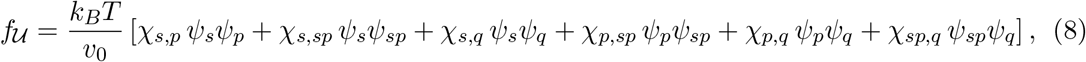

where the Flory–Huggins interaction parameters are

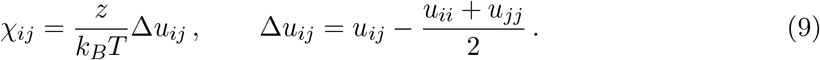

Here, *z* denotes the lattice coordination number, the number of nearest neighbors of a lattice site, and #*u*_*ij*_ is the exchange-energy difference for the *i*–*j* pair. For the four species, the six Flory–Huggins interaction parameters χ_*ij*_ (Eq. 9) are determined by ten distinct interaction energies *u*_*ij*_: four self-interaction and six cross-interaction energies.

#### Energetics of reaction (ℰ)

The reaction-energy contribution in Eq. 6 is

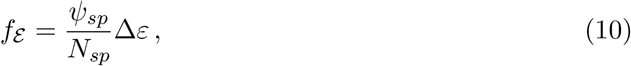

where Δ*ε* is the reaction-energy difference (products minus reactants), and 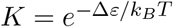 is the corresponding equilibrium association constant. In the ideal case (χ_*ij*_ = 0), the equilibrium relation between the free-species volume fractions is

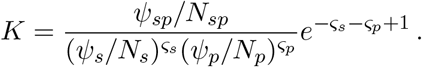

### 2.3 Enthalpy and reaction affinity

For the equilibrium Flory–Huggins calculations, we express the free-species fractions in terms of the total protein and complex fractions,

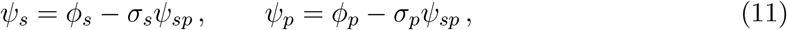

where *σ*_*s*_, *σ*_*p*_, and *N*_*sp*_ are defined above. At fixed *ϕ*_*s*_ and *ϕ*_*p*_, the admissible homogeneous stationary complex fractions satisfy

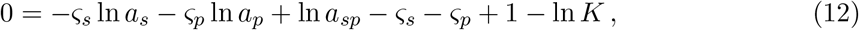

or equivalently,

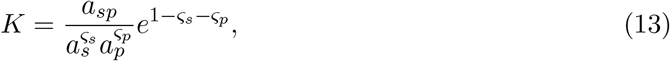

where the molecular activities *a*_*i*_ and activity coefficients *γ*_*i*_ are

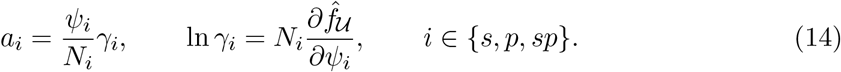

Here, 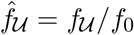 with *f*_0_ = *k*_*B*_ *T/v*_0_. Combining Eqs. 11–14 gives

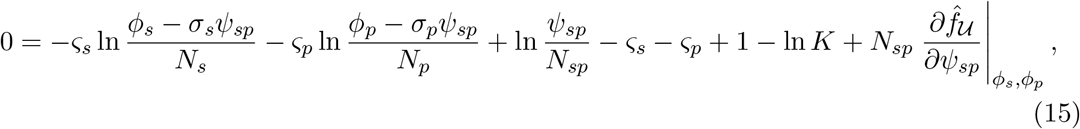

where

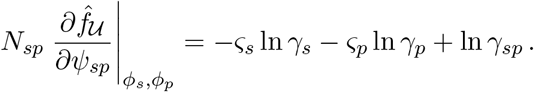

The constrained enthalpic derivative depends on *ψ*_*sp*_, *ϕ*_*s*_, and *ϕ*_*p*_,

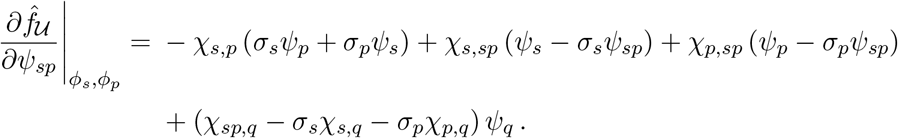

At each total composition, the *ψ*_*sp*_(*ϕ*_*s*_, *ϕ*_*p*_) solving Eq. 15 denote the admissible stationary root(s) that minimize *f*_bulk_. Appendix G describes the selection procedure when multiple roots exist.

The Flory–Huggins free energy described above provides the equilibrium framework for calculating binodals, smooth-branch spinodals, and association-branch switch loci for SynGAP–PSD-95 LLPS (see §3).

### 2.4 Cahn–Hilliard–reaction simulations

For Cahn–Hilliard–reaction simulations, we do not impose local reaction equilibrium. Variational derivatives of Eq. 2 yield chemical potentials conjugate to the species volume fractions,

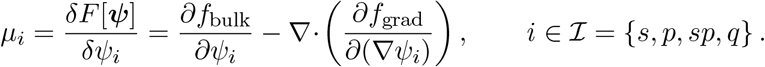

The four-species Cahn–Hilliard–reaction system is

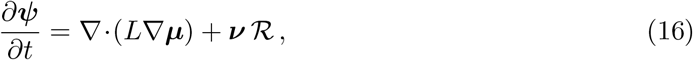

where ***ψ*** = (*ψ*_*s*_, *ψ*_*p*_, *ψ*_*sp*_, *ψ*_*q*_)^⊤^, ***µ*** = (*µ*_*s*_, *µ*_*p*_, *µ*_*sp*_, *µ*_*q*_)^⊤^, and ***ν*** = (−*σ*_*s*_, −*σ*_*p*_, 1, 0)^⊤^.

ℛ denotes the reaction contribution to the complex volume-fraction rate, with ℛ > 0 for net association. We use explicit forward and reverse activity-based mass-action rates

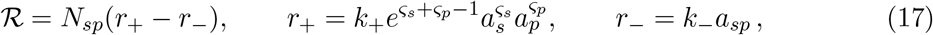

where *k*_+_ and *k*_−_ are association and dissociation rate constants, respectively, with physical dimensions of 1*/*T. Appendix B discusses the relationship between these rates and corresponding rate constants for species concentrations.

For pairwise exchange mobilities *M*_*i,j*_ = *M*_*j,i*_ *≥* 0 for *i ≠ j*, the symmetric Onsager mobility matrix *L* is defined by

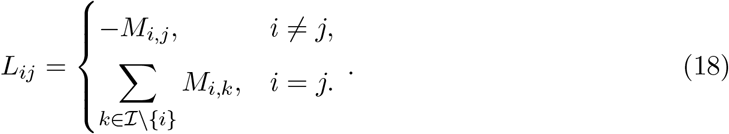

The pairwise mobilities are constructed from nonnegative species mobility weights *m*_*i*_ according to

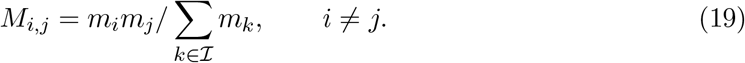

Substituting the pairwise-mobility form of *L* into Eq. 16 gives, for each *i* ∈ ℐ,

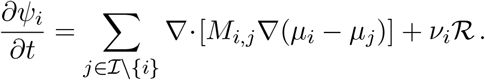

Because local incompressibility imposes *ψ*_*q*_ = 1 − *ψ*_*s*_ − *ψ*_*p*_ − *ψ*_*sp*_, we eliminate the solvent field and define the solvent-referenced chemical-potential differences

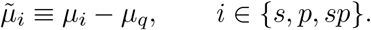

Equivalently, the 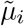 are variational derivatives of the reduced free-energy functional obtained after substituting *ψ*_*q*_ = 1 − *ψ*_*s*_ − *ψ*_*p*_ − *ψ*_*sp*_. We define

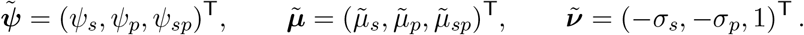

The reduced mobility matrix is the leading principal block 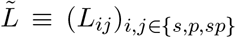 ^. Let **1** =^ (1, 1, 1, 1)^⊤^. Because *L***1** = **0**, *L∇* ***µ*** = *L ∇* (***µ*** − *µ*_*q*_**1**), the first three rows of Eq. 16 are

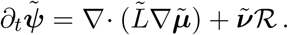

In the Cahn–Hilliard–reaction simulations presented below,

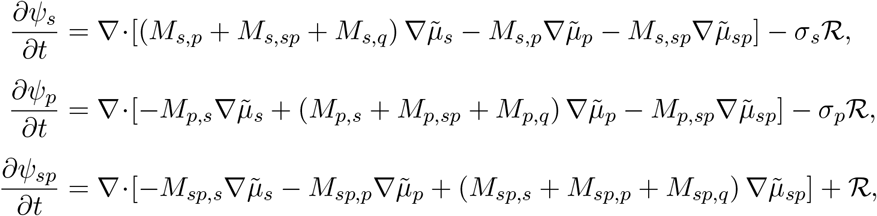

with *ψ*_*q*_ = 1 − *ψ*_*s*_ − *ψ*_*p*_ − *ψ*_*sp*_. Appendix A describes the physical dimensions of the model quantities and the nondimensionalization used in the simulations.

### 2.5 Further modeling assumptions

#### Association of SynGAP trimers and PSD-95

As specified in Eq. 4, free SynGAP (*S*) and PSD-95 (*P*) associate reversibly. For stoichiometric coefficients *ς*_*s*_ = 3 and *ς*_*p*_ = 2, Eq. 4 becomes

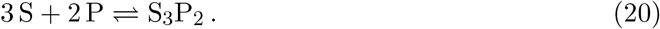

Because SynGAP exists predominantly as a trimer, we follow Lin *et al*. [17] and rewrite Eq. 20 in terms of SynGAP trimers as

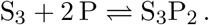

In this trimer-based scheme, the SynGAP stoichiometric coefficient is 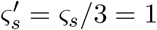, and 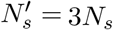 is the trimer relative-volume parameter. This reformulation leaves the SynGAP volume fraction within the complex, *σ*_*s*_ = *ς*_*s*_*N*_*s*_*/N*_*sp*_, unchanged because 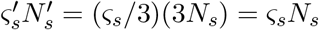. Hereafter, we adopt this trimer-based representation and suppress the primes: *S* denotes one SynGAP trimer with *ς*_*s*_ = 1, and *N*_*s*_ is the relative molecular volume of the trimer.

The relative-volume parameters *N*_*i*_ are effective Flory–Huggins size parameters. We take PSD-95 and one coarse-grained solvent/background unit as reference sizes, setting *N*_*p*_ = *N*_*q*_ = 1; thus, *Q* represents an effective aqueous background volume rather than an individual solvent molecule. The physically motivated construct-like and full-length-like calibrations use *N*_*s*_ = 1 and *N*_*s*_ = 5, respectively; selected equilibrium sensitivity calculations additionally use *N*_*s*_ = 3 as an illustrative intermediate value rather than as a separate physical calibration (see Appendix A).

#### Volume fractions *ψ*_*i*_ and association constant *K*

Fig. 4 shows how the equilibrium association constant *K* affects the species volume fractions *ψ*_*s*_ (free SynGAP), *ψ*_*p*_ (free PSD-95), *ψ*_*sp*_ (SynGAP–PSD-95 complex), and *ψ*_*q*_ (solvent) when there is no enthalpic contribution to the bulk free energy (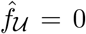 in Eq. 6). In the limiting case *K* = 0, no complex forms (*ψ*_*sp*_ = 0), so the free-species fractions equal the total fractions: *ψ*_*i*_ = *ϕ*_*i*_ for *i* ∈ *{s, p, q}* (Eq. 5) and the ternary contours are exactly the barycentric coordinate fields (Fig. 4**A**). Fig. 4**B** and Fig. 4**C** show weakly (*K* = 10) and strongly (*K* = 10^3^) favorable association between *S* and *P*, respectively, for (*N*_*s*_, *N*_*p*_, *N*_*q*_) = (1, 1, 1). Fig. 4**D** repeats the strongly favorable case for (*N*_*s*_, *N*_*p*_, *N*_*q*_) = (5, 1, 1). For the favorable-association cases, the complex fraction *ψ*_*sp*_ is smallest when either total protein fraction approaches zero and largest at intermediate values of both *ϕ*_*s*_ and *ϕ*_*p*_. When χ_*ij*_ = 0 and *ϕ*_*q*_ is fixed, the total composition 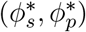 that maximizes *ψ*_*sp*_ satisfies the free-species ratio

**Figure 4.**
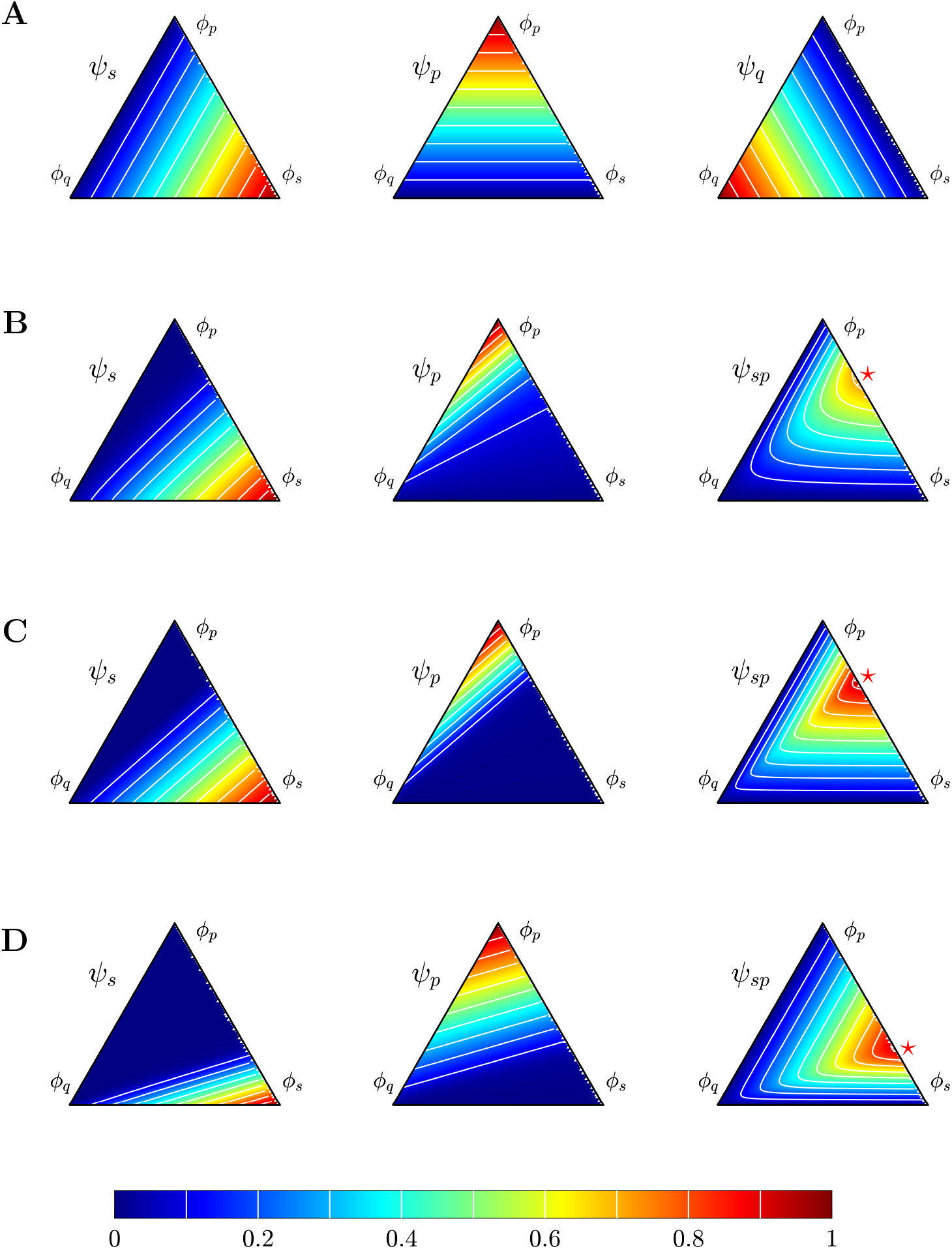
Effiect of the equilibrium association constant *K* on species volume fractions. Ternary diagrams are parameterized by the total volume fractions ***ϕ*** = (*ϕ*_*s*_, *ϕ*_*p*_, *ϕ*_*q*_), constrained by *ϕ*_*s*_+*ϕ*_*p*_+*ϕ*_*q*_ = 1, and use complex stoichiometry (*ς*_*s*_, *ς*_*p*_) = (1, 2). All interaction energies are zero, isolating the effect of SynGAP–PSD-95 association (Eq. 4). Pseudocolor shows the indicated species volume fraction, with white contours at 0.1 intervals from 0.1 through 0.9. Panels **A–C** use (*N*_*s*_, *N*_*p*_, *N*_*q*_) = (1, 1, 1). **A**: *ψ*_*s*_, *ψ*_*p*_, and *ψ*_*q*_ in the limiting case *K* = 0. **B,C**: *ψ*_*s*_, *ψ*_*p*_, and *ψ*_*sp*_ at *K* = 10 and 10^3^, respectively. **D**: *ψ*_*s*_, *ψ*_*p*_, and *ψ*_*sp*_ at *K* = 10^3^ with (*N*_*s*_, *N*_*p*_, *N*_*q*_) = (5, 1, 1). Red stars in **B–D** mark the corresponding solvent-free stoichiometric compositions at which *ψ*_*sp*_ is maximal in the high-affinity limit (Eq. 21).

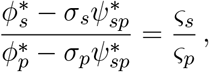

where 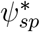 denotes the maximum complex fraction. In the high-affinity limit *K → ∞, ψ*_*sp*_ approaches the limiting-reactant bound from below according to

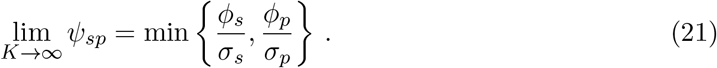

In this limit, the maximum 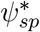 occurs at 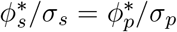, where neither SynGAP nor PSD-95 is in stoichiometric excess. Increasing *K* promotes complex formation, increasing *ψ*_*sp*_ while decreasing the free-protein fractions *ψ*_*s*_ and *ψ*_*p*_. Accordingly, *ψ*_*sp*_ is largest along the solvent-free edge (*ψ*_*q*_ = 0) near the stoichiometric ratio *ϕ*_*s*_*/ϕ*_*p*_ = *σ*_*s*_*/σ*_*p*_. This ratio is 1*/*2 for (*N*_*s*_, *N*_*p*_, *N*_*q*_) = (1, 1, 1) in Fig. 4**B,C** and 5*/*2 for (5, 1, 1) in Fig. 4**D**; red ⋆ symbols mark these maxima.

#### Flory–Huggins interaction energetics

For four species, the Flory–Huggins enthalpy (Eq. 8) contains six dimensionless interaction parameters χ_*ij*_, obtained from ten interaction energies *u*_*ij*_. For simplicity, interactions involving the SynGAP–PSD-95 complex are defined as additive sums of the corresponding constituent interactions:

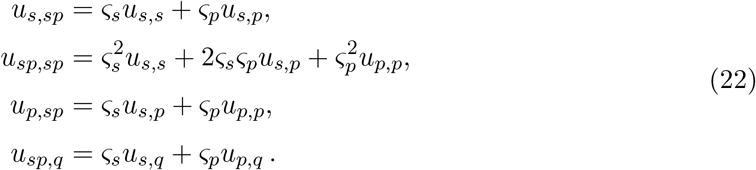

Appendix C details how substitution into Eq. 9 gives the map from the six constituent interaction energies (*u*_*ij*_) to six exchange-energy differences (Δ*u*_*ij*_) and six Flory–Huggins interaction parameters (χ_*ij*_ = *z*Δ*u*_*ij*_*/k*_*B*_*T*).

## 3 Results

### 3.1 Ternary Flory–Huggins diagram showing complex-mediated phase separation

To characterize SynGAP–PSD-95 complex-mediated phase separation, we numerically evaluated the equilibrium free energy across the total-composition simplex and identified binodals, smooth-branch spinodals, and association-branch switch loci for multiple values of *K*. Fig. 5 shows the resulting Flory–Huggins free-energy density *f*_bulk_ (Eq. 6) as a function of the total volume fractions ***ϕ***. We approximate the equilibrium mixture free-energy density using the lower convex envelope of *f*_bulk_ on the composition mesh; compositions at which *f*_bulk_ exceeds this envelope lie in the phase-coexistence region (see Appendix F for details). At the displayed total compositions, we verify the lower-hull construction by separately minimizing the free energy stochastically over lever-rule-compatible phase fractions and compositions (Appendix F.1). Standard parameters are listed in Table 1, and the figure-specific values of *K* and the elementary interaction energies *u*_*i,j*_ are stated in the caption.

**Table 1.** Standard model parameters and auxiliary definitions. Unless otherwise stated, the standard parameters are fixed across each pre-/post-LTP comparison; condition-specific thermodynamic parameters are listed in Table 2. Figure-specific total compositions are stated in the corresponding figure captions.

| Symbol | Definition | Standard value or relation |
| --- | --- | --- |
| **Standard parameters** |  |  |
| $(N_s, N_p, N_q)$ | Relative molecular volumes | $(3, 1, 1)$ |
| $(\varsigma_s, \varsigma_p)$ | Complex stoichiometry | $(1, 2)$ |
| $(u_{s,s}, u_{s,p}, u_{s,q}, u_{p,p}, u_{p,q})$ | Constituent interaction energies | varied |
| $(\phi_s^0, \phi_p^0, \phi_q^0)$ | Total composition | varied |
| $K$ | Equilibrium association constant | varied |
| $k_+$ | Forward mass-action rate constant | Appendix A |
| $\kappa_i$ | Square-gradient coefficients | Appendix A |
| $(m_s, m_p, m_{sp}, m_q)$ | Species mobility weights | Appendix A |
| **Auxiliary definitions** |  |  |
| $N_{sp}$ | Relative molecular volume of complex | $N_{sp} = \varsigma_s N_s + \varsigma_p N_p$ |
| $(\sigma_s, \sigma_p)$ | Complex-volume fractions | $(\sigma_s, \sigma_p) = (\varsigma_s N_s / N_{sp}, \varsigma_p N_p / N_{sp})$ |
| $(u_{s,sp}, u_{p,sp}, u_{sp,sp}, u_{sp,q})$ | Composite interaction energies | Eq. 22 |
| $\Delta u_{ij}$ | Exchange-energy difference | $\Delta u_{ij} = u_{ij} - \frac{1}{2}(u_{ii} + u_{jj})$ |
| $\chi_{ij}$ | Flory–Huggins parameters | $\chi_{ij} = (z/k_B T) \Delta u_{ij}; z = 6$ |
| $\Delta \varepsilon$ | Reaction-energy difference | $\Delta \varepsilon = -k_B T \ln K$ |
| $k_-$ | Reverse mass-action rate constant | $k_- = k_+ / K$ |
| $L_{ij}$ | Onsager mobilities | Eq. 18 |
| $M_{i,j}$ | Pairwise mobilities | Eq. 19 |

**Figure 5.**
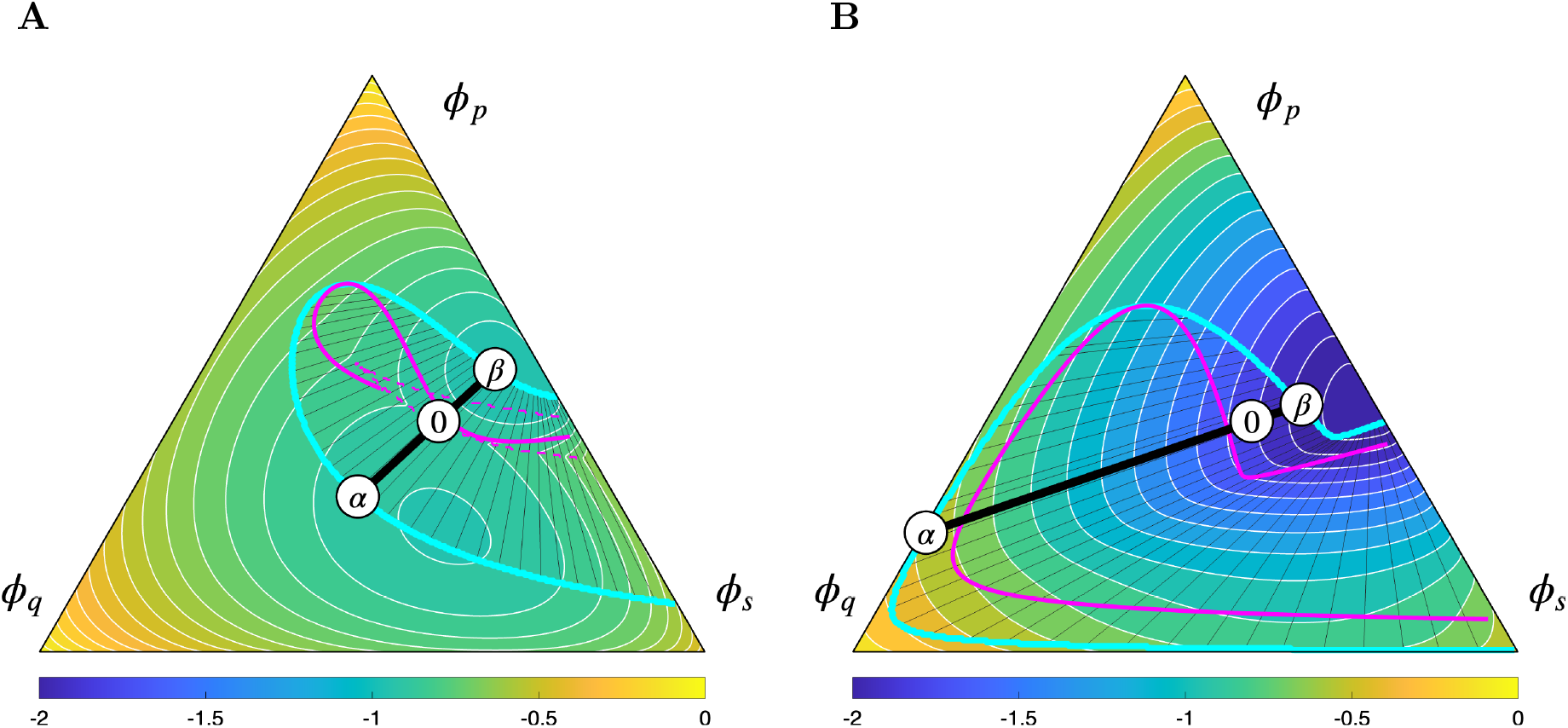
Equilibrium association constant *K* and phase behavior. Ternary Flory– Huggins phase diagrams in total-composition space, calculated on a 301-point-per-axis simplex mesh. Pseudocolor shows the homogeneous bulk free-energy density *f*_bulk_. In each panel, the thick black tie line connects the coexisting *α* and *γ* compositions through the representative total composition ***ϕ***^0^ = (0.4, 0.4, 0.2), indicated by the white circle labeled 0. Thin black lower-hull edges show a representative subset of the surrounding tie-line structure. Cyan curves mark the binodals. Solid magenta curves mark smooth-branch spinodals; in **A**, dashed magenta curves mark the global-minimizer branch-switch locus at which spinodal segments terminate. **A**: Phase behavior at *K* = 10. **B**: Phase behavior at *K* = 10^4^. Both panels use *u*_*ss*_ = *u*_*sp*_ = 0, and *u*_*pp*_ = −1*/*6 with solvent interaction energies set to zero. Both panels use the illustrative intermediate relative-volume choice (*N*_*s*_, *N*_*p*_, *N*_*q*_) = (3, 1, 1), which is distinct from the construct-like (1, 1, 1) and full-length-like (5, 1, 1) physical calibrations (Appendix A). Remaining standard parameters are listed in Table 1.

For the two-phase calculations beginning with Fig. 5, we label the coexisting phases with lower and higher total PSD-95 volume fraction *ϕ*_*p*_ as *α* and *β*, respectively. These labels denote the PSD-95-poor and PSD-95-rich phases and do not imply an ordering by total solute or solvent content. At *K* = 10, Fig. 5A shows a relatively small two-phase region concentrated near compositions with the largest equilibrium complex fraction (cf. Fig. 4). At fixed interaction parameters, increasing the association constant to *K* = 10^4^ greatly expands the two-phase region (Fig. 5B). For compositions inside the two-phase region at *K* = 10^4^ but outside it at *K* = 10, increasing *K* between these values changes the equilibrium from homogeneous to two-phase; reversing the change restores homogeneity.

### 3.2 Effect of interaction parameters on phase separation

We next examine how varying the selfand cross-interaction energies *u*_*ss*_, *u*_*pp*_, and *u*_*sp*_ affects phase separation at fixed equilibrium association constant *K* = 10^4^ (Fig. 6).

**Figure 6.**
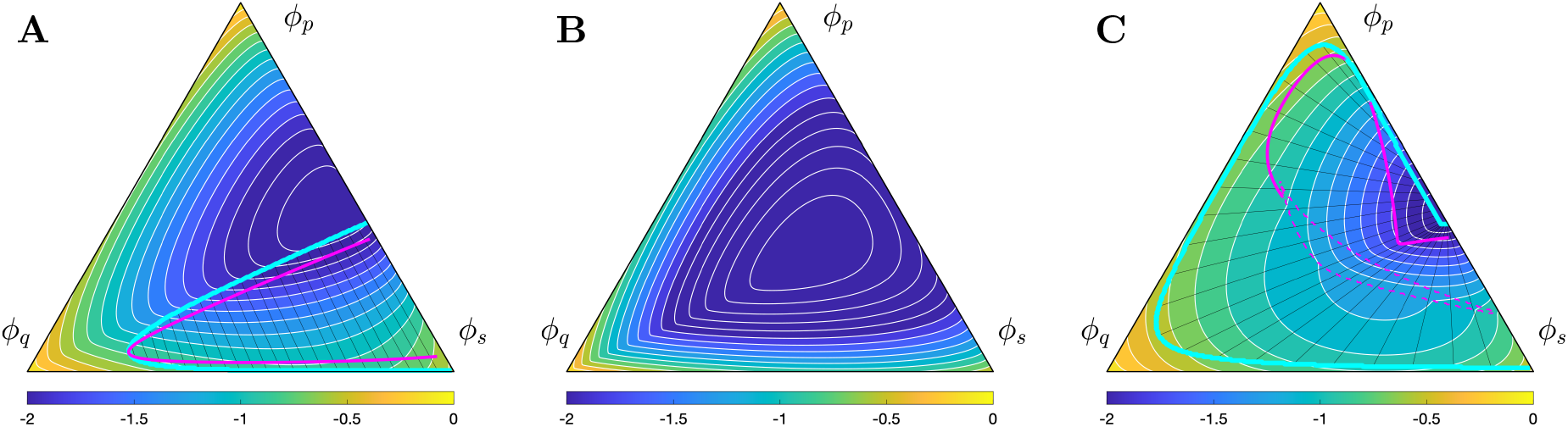
Effiect of interaction energies on phase behavior. Ternary Flory–Huggins phase diagrams in total-composition space for the four-species system (*ψ*_*s*_, *ψ*_*p*_, *ψ*_*sp*_, *ψ*_*q*_), obtained by varying *u*_*ij*_ at fixed equilibrium association constant *K* = 10^4^. Pseudocolor shows the homogeneous bulk free-energy density *f*_bulk_. Annotations and phase boundary conventions are as in Fig. 5. **A**: Reference interactions, *u*_*s,s*_ = *u*_*s,p*_ = 1*/*6 and *u*_*p,p*_ = −1*/*6. **B**: Repulsive PSD-95–PSD-95 interactions, *u*_*p,p*_ = 1*/*6. **C**: Attractive SynGAP–PSD-95 interactions, *u*_*s,p*_ = −1*/*6. Remaining parameters are as in Fig. 5.

Fig. 6A shows the reference phase diagram, whereas panels B and C show the effects of changing *u*_*pp*_ and *u*_*sp*_, respectively. Making the PSD-95–PSD-95 interaction repulsive (*u*_*pp*_ = 1*/*6; Fig. 6B) nearly eliminated the two-phase region. Making the SynGAP–PSD-95 interaction attractive (*u*_*sp*_ = −1*/*6; Fig. 6C) substantially expanded the two-phase region. Changing *u*_*ss*_ from 1*/*6 to −1*/*6 while holding the other parameters fixed made the SynGAP– SynGAP interaction attractive but produced no discernible change relative to the reference case (not shown).

Fig. 8 shows that the equilibrium association constant and relative molecular volumes jointly shape the phase diagram. For each of the three displayed relative-volume choices, increasing *K* from 1 to 1000 progressively expands both the two-phase and locally unstable regions. At fixed *K* = 100 or 1000, increasing *N*_*s*_ from 1 to 5 reduces the extent of both regions, whereas the small two-phase region near the onset at *K* = 1 depends nonmonotonically on *N*_*s*_. Thus, the qualitative promotion of phase separation by stronger association is robust across the tested relative volumes, but the locations and extents of the phase boundaries are not.

**Figure 7.**
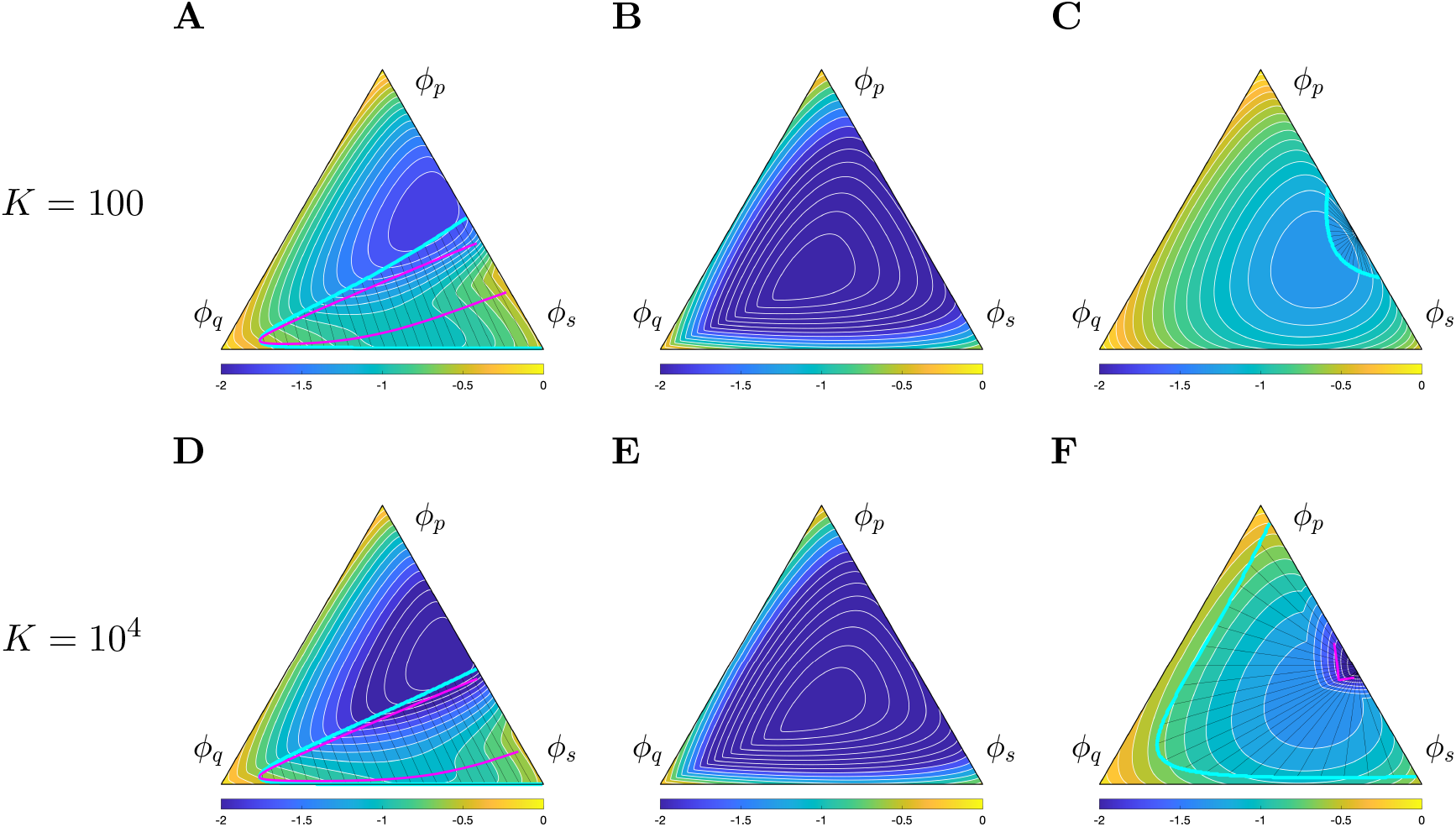
Combined effiects of *K* and interaction energies on phase behavior. Ternary Flory–Huggins phase diagrams in total-composition space for the four-species system (*ψ*_*s*_, *ψ*_*p*_, *ψ*_*sp*_, *ψ*_*q*_), by varying both *K* and the interaction energies *u*_*ij*_. Pseudocolor, annotations, and phase boundary conventions are as in Fig. 5. Panels **A–C** use *K* = 100, whereas panels **D–F** use *K* = 10^4^. Within each group, panels **A** and **D** show the reference interaction energies (*u*_*s,s*_ = *u*_*s,p*_ = 1*/*3, *u*_*p,p*_ = −1*/*3); panels **B** and **E** use repulsive PSD95–PSD-95 interactions (*u*_*p,p*_ = 1*/*3); and panels **C** and **F** use attractive SynGAP–PSD-95 interactions (*u*_*s,p*_ = −1*/*3). Remaining parameters are as in Fig. 5.

**Figure 8.**
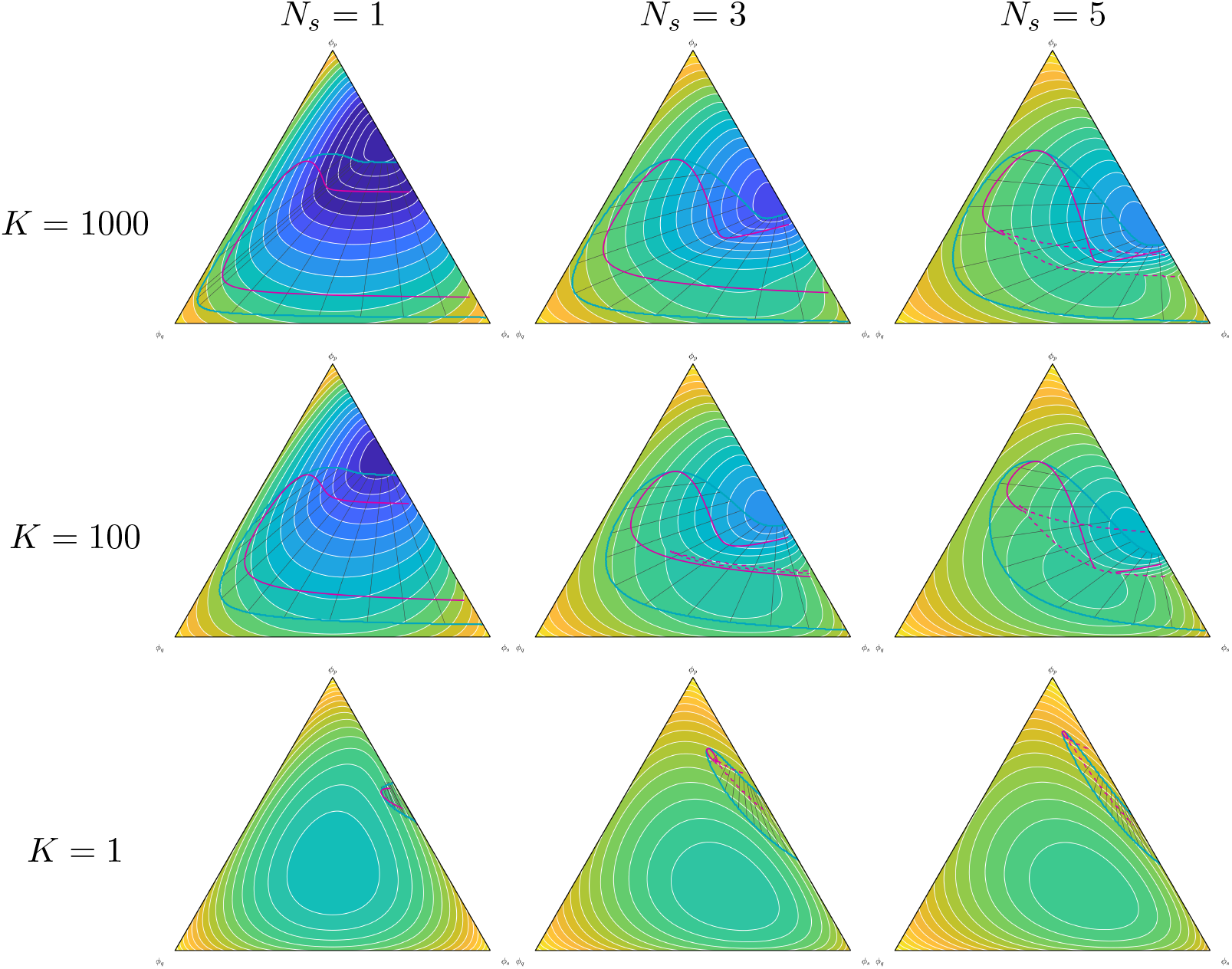
Relative-volume and association-strength sensitivity of the two-phase region. The equilibrium calculations underlying Fig. 5 were repeated at (*N*_*s*_, *N*_*p*_, *N*_*q*_) = (1, 1, 1), (3, 1, 1), and (5, 1, 1) (columns) for association constants *K* = 1000, 100, and 1 (rows). All interaction energies and other numerical parameters are unchanged. Pseudocolor, annotation, and phase boundary conventions are as in Fig. 5.

### 3.3 Illustrative LTP-like parameter shift

Because equilibrium phase behavior depends on both complex formation and intermolecular interaction energies, we examine how an illustrative LTP-like parameter switch changes the computed two-phase region and coexisting phase compositions. During chemically induced LTP, SynGAP rapidly disperses from dendritic spines and remains dispersed at 60 minutes in stably responding spines. This observation suggests that activity-dependent signaling alters SynGAP partitioning and its effective affinity for PSD-95 [3]. Fig. 9 illustrates this change phenomenologically using pre- and post-LTP parameter sets that differ in *K* and selected interaction energies. The pre- and post-switch parameters are listed in Table 2.

**Table 2.** Pre- and post-LTP thermodynamic parameters for Figs 9–11. Solvent interaction energies are zero (*u*_*s,q*_ = *u*_*p,q*_ = *u*_*sp,q*_ = *u*_*q,q*_ = 0); other fixed parameters and auxiliary definitions are listed in Table 1.

| Symbol | Definition | Pre-LTP | Post-LTP |
| --- | --- | --- | --- |
| $K$ | Equilibrium association constant | $10^3$ | $10^{-3}$ |
| $u_{s,s}$ | SynGAP–SynGAP interaction energy | $1/6$ | $-1/4$ |
| $u_{s,p}$ | SynGAP–PSD-95 interaction energy | $-1/4$ | $1/6$ |
| $u_{p,p}$ | PSD-95–PSD-95 interaction energy | $-1/12$ | $-1/12$ |

**Table 3.**
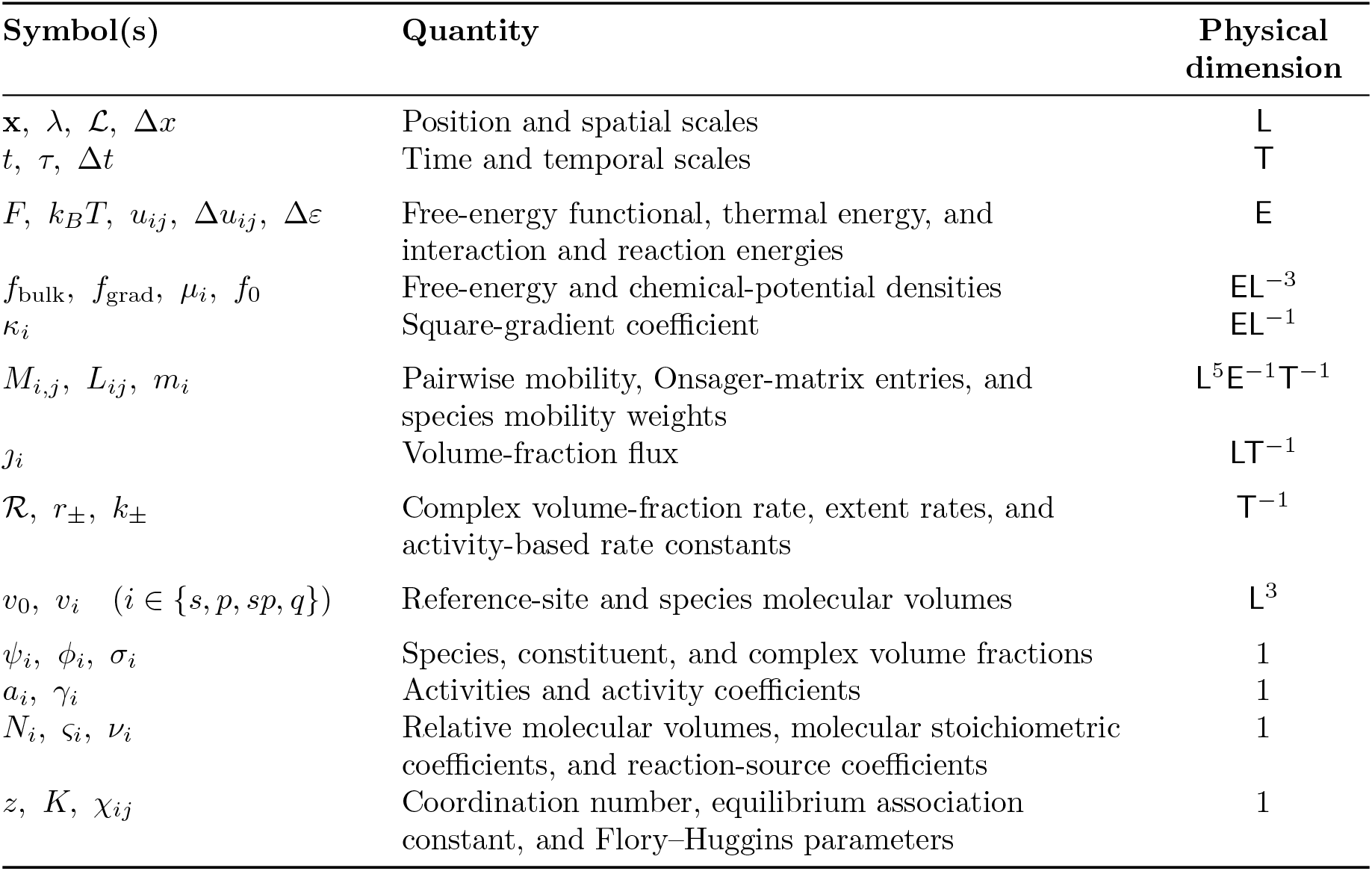
Physical dimensions of the quantities entering the Cahn–Hilliard–reaction model. An entry of 1 denotes a dimensionless quantity.

**Figure 9.**
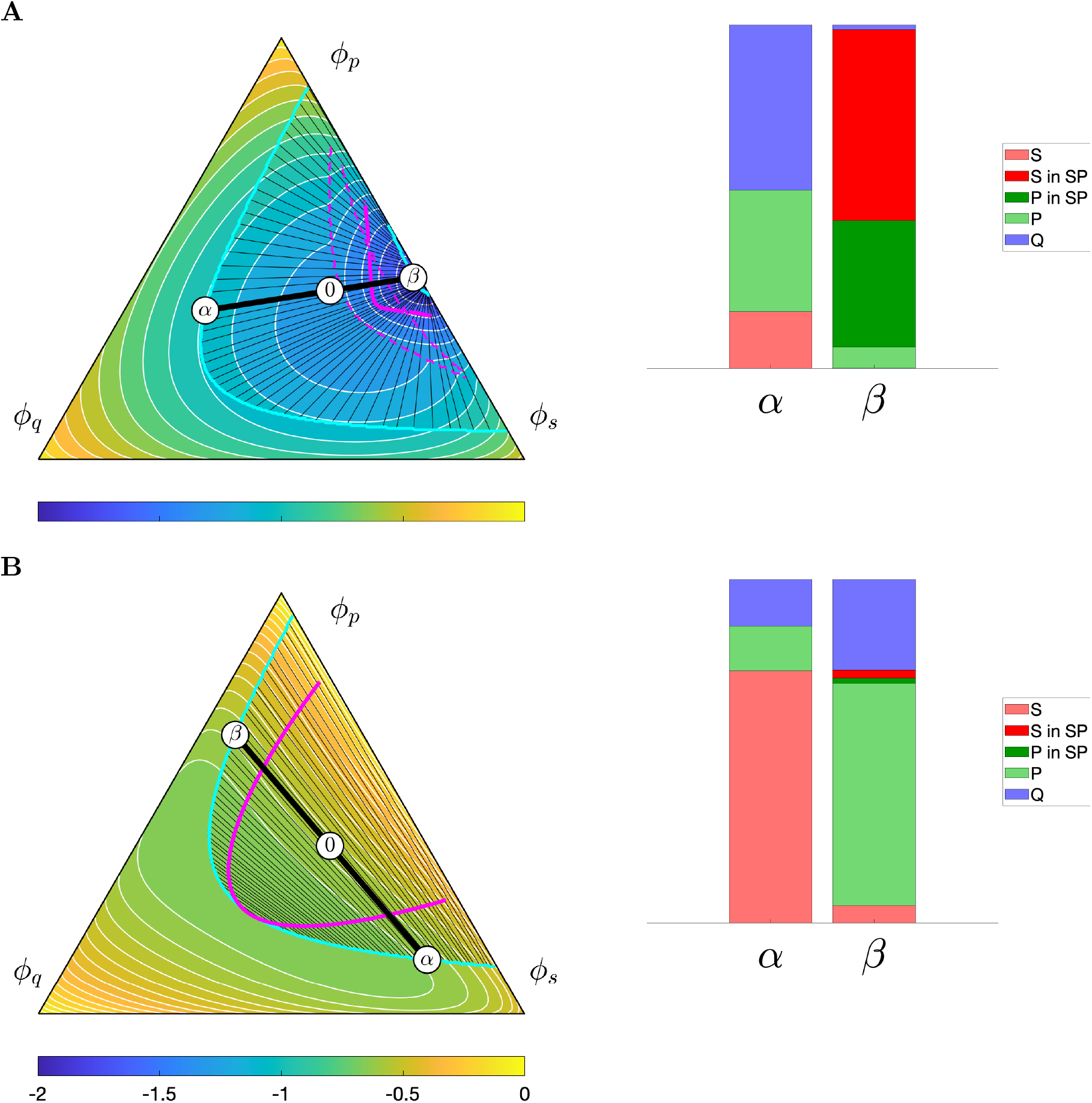
Phase diagrams before and after an illustrative LTP-like parameter switch. Changing *K* and selected *u*_*ij*_ values shifts the tie lines and increases free SynGAP in the *α* phase at the chosen total composition. Pseudocolor, annotations, and phase boundary conventions are as in Fig. 5. The thick black tie line passes through ***ϕ***^0^ = (0.4, 0.4, 0.2) and connects the labeled coexistence compositions. The adjacent bars decompose each phase into free S, S in SP, P in SP, free P, and solvent Q. Here SP denotes the SynGAP–PSD-95 complex (Eq. 4). **A**: Pre-LTP condition. **B**: Post-LTP condition. Condition-specific parameters are listed in Table 2; remaining parameter values are as in Fig. 5.

In each ternary diagram, the thick tie line through ***ϕ***^0^ = (0.4, 0.4, 0.2) identifies the coexisting PSD-95-poor *α* and PSD-95-rich *β* phases, and the adjacent stacked bars resolve those same tie-line endpoints into free proteins, the SynGAP and PSD-95 contributions to complex, and solvent (Fig. 9). Before the switch, the *β* phase is dominated by complex, whereas after the switch the complex contribution is small and free SynGAP and free PSD-95 partition preferentially into the *α* and *β* phases, respectively. The bars therefore show that the shifted tie lines correspond to a redistribution of the molecular species between phases, rather than a loss of two-phase coexistence.

The illustrative pre-LTP parameter set (Fig. 9A) represents strong SynGAP–PSD-95 association with *K* = 10^3^, attractive PSD-95–PSD-95 and SynGAP–PSD-95 interactions (*u*_*pp*_ = −1*/*12, *u*_*sp*_ = −1*/*4), and a repulsive SynGAP–SynGAP interaction (*u*_*ss*_ = 1*/*6). These parameters yield coexistence of a PSD-95-rich phase (*β*) and a PSD-95-poor phase (*α*).

The illustrative post-LTP parameter set reduces the association constant to *K* = 10^−3^ and changes the interaction energies to *u*_*ss*_ = −1*/*4, *u*_*sp*_ = 1*/*6, and *u*_*pp*_ = −1*/*12. Relative to the pre-LTP parameter set, these changes make the SynGAP–PSD-95 interaction less favorable and the SynGAP–SynGAP interaction more favorable, while leaving the PSD-95– PSD-95 interaction unchanged. For this illustrative parameter shift, phase separation persists while SynGAP redistributes from the PSD-95-rich *β* phase to the PSD-95-poor *α* phase.

### 3.4 Post-switch spatial redistribution

We initialize the pre-LTP calculations from coexistence profiles constructed using the equilibrium *α*- and *β*-phase compositions

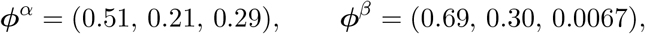

with a lever-rule beta-phase fraction of *v*_*β*_ = 0.10. The mean composition is

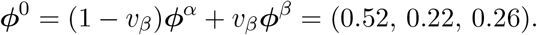

The initial profile consists of a ∼100-nm interval in one dimension and a circular *β*-phase region in two dimensions.

In both one and two spatial dimensions, we initialize the simulation with a pre-LTP profile that is nearly at steady state. At relative post-switch time *t* = 0 (absolute simulation time *t* = 10 s), the association constant *K* is decreased from 10^3^ to 10^−3^ and the interaction energies are switched to their post-LTP values. The simulation continues for 60 s after the switch. We visualize these dynamics using sequential one- and two-dimensional snapshots of the total and species-resolved compositions at relative times 1, 3, 10, 30, and 60 s (Fig. 10).

**Figure 10.**
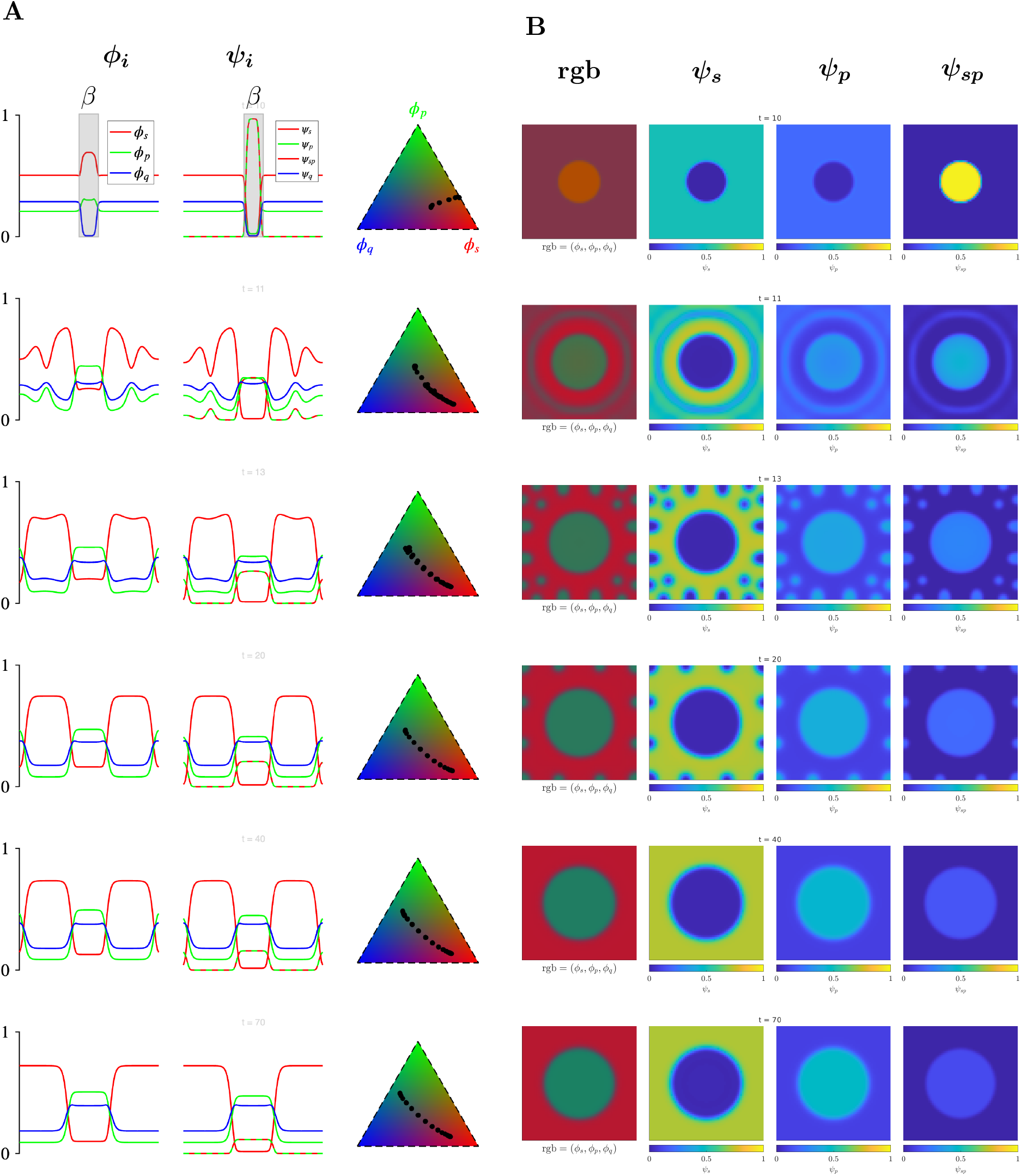
Cahn–Hilliard–reaction dynamics following the illustrative LTP-like parameter switch. Pre-LTP simulations are initialized from the corresponding equilibrium *α*and *β*-phase compositions with beta-phase fraction 0.10, using a periodic box profile in one dimension and a circular *β*-phase region in two dimensions. In both panels, the state after 10 s of pre-LTP evolution is restarted using the post-LTP parameters; rows show absolute times *t* = 10, 11, 13, 20, 40, and 70 s, corresponding to relative post-switch times 0, 1, 3, 10, 30, and 60 s. **A**: In the first row, the *β*-phase is marked with a gray rectangle. One-dimensional calculation with 128 grid points (Δ*x* = 7.8125 nm). **B**: Two-dimensional calculation with a 64 × 64 grid (Δ*x* = 15.625 nm). In each panel, the first row is both the selected *before* checkpoint and the initial *after* state recorded before the first post-switch time step, confirming restart continuity. Both simulations use a 1000-nm periodic domain, nominal time step Δ*t* = 10^−4^ s, 16-nm gradient length, and mean total composition ***ϕ***^0^ = (0.5238, 0.2190, 0.2572). The provisional collective diffusivities for (*S, P, SP, Q*) are (0.012, 0.020, 0.010, 0.020) *µ*m^2^ s^−1^, and the post-switch dissociation prefactor corresponds to a 60-s timescale.

Fig. 10A shows the one-dimensional evolution; the left column gives periodic phase-separated profiles of the total volume fractions *ϕ*_*s*_, *ϕ*_*p*_, and *ϕ*_*q*_. In the pre-LTP total-composition profiles, *ϕ*_*s*_ (red) and *ϕ*_*p*_ (green) vary in phase and both vary out of phase with the solvent fraction *ϕ*_*q*_, so protein-rich peaks align with solvent-fraction troughs. The species-resolved profiles (*ψ*_*s*_, *ψ*_*p*_, *ψ*_*sp*_, *ψ*_*q*_) show the free SynGAP, free PSD-95, complex, and solvent contributions separately. Before LTP, the complex fraction *ψ*_*sp*_ varies in phase with the free PSD-95 fraction *ψ*_*p*_, and both are depleted where the free SynGAP fraction *ψ*_*s*_ peaks.

In the post-LTP total-composition profiles, *ϕ*_*s*_ (red) and *ϕ*_*p*_ (green) vary out of phase, whereas *ϕ*_*p*_ and the solvent fraction *ϕ*_*q*_ (blue) vary in phase. After the parameter switch, *ψ*_*sp*_ decreases across the displayed times while remaining in phase with *ψ*_*p*_ and out of phase with *ψ*_*s*_. Its reduction by the final displayed snapshot is consistent with complex dissociation while spatial phase separation persists (cf. Fig. 9).

Fig. 10B shows the corresponding two-dimensional evolution. The left column shows RGB composites that map the total fractions (*ϕ*_*s*_, *ϕ*_*p*_, *ϕ*_*q*_) to the red, green, and blue channels, respectively. Early amber regions indicate overlap of total SynGAP and PSD-95, whereas purple regions indicate overlap of total SynGAP and solvent. After the parameter switch, the RGB composites evolve toward red–green partitioning of the total SynGAP and PSD-95 fractions. Secondary composition domains appear in the material surrounding the central domain by relative time 3 s but subsequently recede. By relative time 30 s, the displayed field has returned to a single central domain, which persists through the final snapshot at 60 s.

The three right columns show the species-resolved fields *ψ*_*s*_, *ψ*_*p*_, and *ψ*_*sp*_ throughout the transition. Before the parameter switch, *ψ*_*sp*_ is strongly enriched in the PSD-95-rich domains, whereas free SynGAP *ψ*_*s*_ is depleted there. At this stage, free PSD-95 *ψ*_*p*_ is also low, and most protein in these domains is complex. Because the complex contributes to both total SynGAP *ϕ*_*s*_ and total PSD-95 *ϕ*_*p*_, these domains appear as mixed colors in the RGB composite. After the parameter switch, *ψ*_*sp*_ decreases over the displayed times, while free SynGAP and free PSD-95 become increasingly enriched in complementary regions.

### 3.5 Haploinsufficiency

Fig. 11 compares the control composition ***ϕ***^0^ = (0.4, 0.4, 0.2) of Fig. 9 with the SynGAP-haploinsufficient composition ***ϕ***^0^ = (0.2, 0.4, 0.4) for the illustrative pre-LTP and post-LTP parameter sets. This SynGAP-haploinsufficient composition reduces 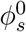 by 50% at fixed 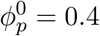, with the remaining volume fraction assigned to solvent.

**Figure 11.**
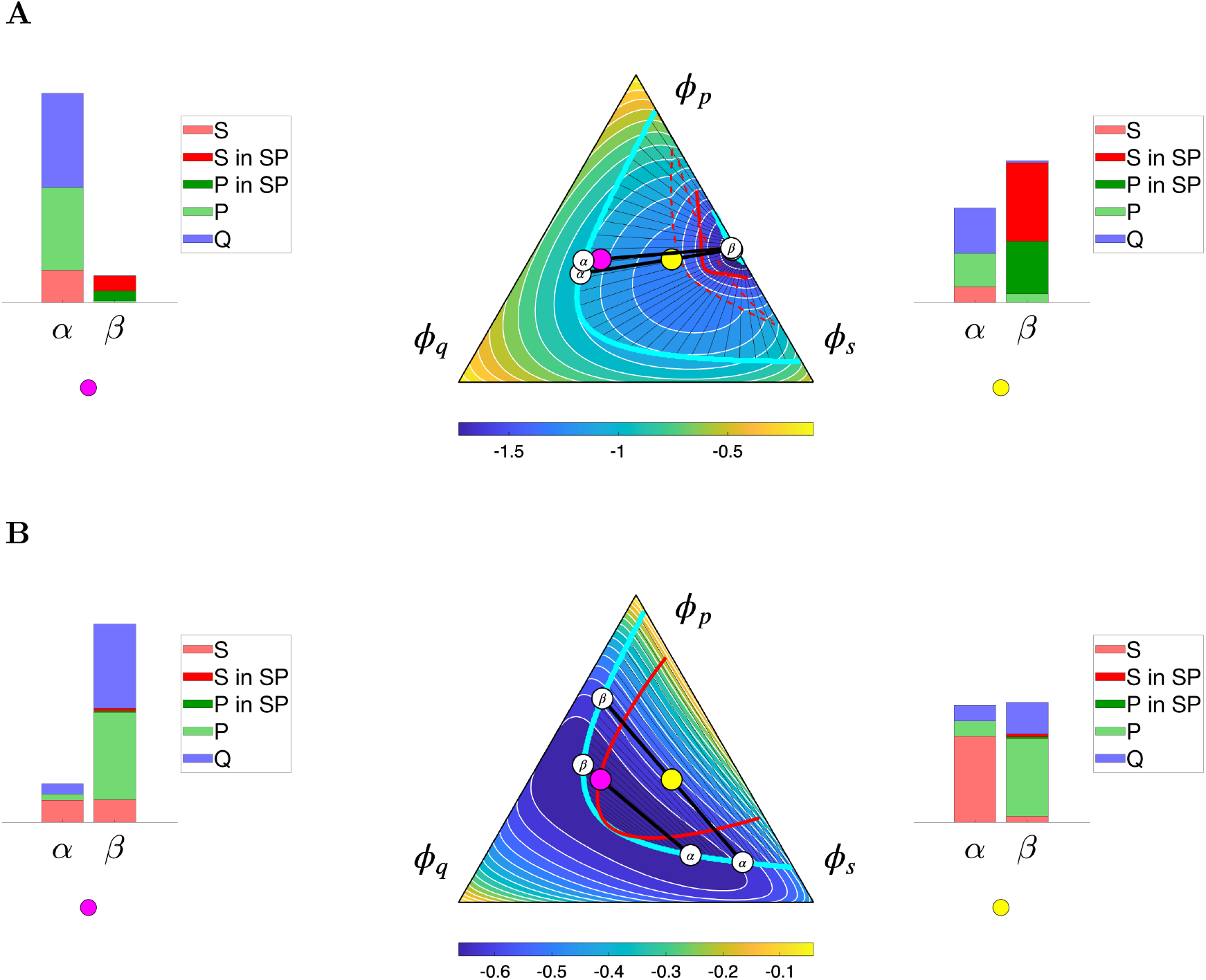
Equilibrium phase behavior at control and reduced SynGAP dosage under illustrative preand post-LTP parameter sets. **A**: Pre-LTP parameters. **B**: Post-LTP parameters. Filled circles identify the SynGAP-haploinsufficient (magenta) and control (yellow) total compositions, ***ϕ***^0^ = (0.2, 0.4, 0.4) and (0.4, 0.4, 0.2), respectively. The stacked bars show the phase-fraction-weighted species contributions for the SynGAP-haploinsufficient composition at left and the control composition at right. The total heights of the *α* and *β* bars equal the lever-rule phase fractions *v*_*α*_ and *v*_*β*_, while the relative colored segments within each bar show that phase’s species composition.

For both parameter sets, the control and SynGAP-haploinsufficient compositions lie within the computed two-phase region. In the pre-LTP row (Fig. 11A), the *β* phase is complex-rich at both dosages. In the post-LTP row (Fig. 11B), the complex contribution is small, and free SynGAP and free PSD-95 instead partition preferentially into the *α* And *β* phases, respectively. Comparing the haploinsufficient bar graph at the left of each row with the corresponding control bar graph at the right, the pre-LTP coexistence compositions change only modestly: the *β* phase remains complex-rich, while the *α* phase contains slightly less free SynGAP and slightly more free PSD-95. In the post-LTP condition, haploinsufficiency increases the solvent content of both phases, reduces the enrichment of free SynGAP in the *α* phase and free PSD-95 in the *β* phase, and further decreases the already small complex contribution. Reduced SynGAP therefore weakens the post-switch protein-partitioning contrast without reversing the preferential localization of SynGAP and PSD-95 to opposing phases. The lever-rule-weighted bar heights additionally show that haploinsufficiency increases the *α*-phase fraction before the switch but increases the *β*-phase fraction after the switch.

## 4 Discussion

We couple a Flory–Huggins free energy in total-composition space to four-species Cahn– Hilliard–reaction dynamics for an idealized SynGAP–PSD-95 system. Explicitly representing the reversible association of one SynGAP trimer with two PSD-95 molecules and the constituent self- and cross-interaction energies links equilibrium phase diagrams to the spatial redistribution of free SynGAP, free PSD-95, their complex, and solvent. Within the model, the equilibrium association constant *K* controls the complex fraction and, together with the interaction energies *u*_*ij*_, determines the extent of the two-phase region [cf. 17]. The effects of the interaction energies depend on the association regime: repulsive PSD-95–PSD-95 interactions reduce the two-phase region, whereas attractive SynGAP–PSD-95 interactions reduce it at *K* = 100 but expand it at *K* = 10^4^ (Figs. 6 and 7).

We represent the LTP-like perturbation phenomenologically by changing *K* and selected interaction energies rather than modeling CaMKII phosphorylation reactions explicitly. The parameter set shown in Fig. 9 illustrates SynGAP redistribution from the PSD-95-rich phase (*β*) into the PSD-95-poor phase (*α*). The fixed total composition remains within the post-switch two-phase region, and a distinct PSD-95-rich phase persists (Fig. 9). This qualitative redistribution is consistent with experimental observations of rapid SynGAP dispersion from synaptic spines during LTP and of its movement out of the postsynaptic-density core upon depolarization [3, 26]. In the spatial simulations, a decrease in *ψ*_*sp*_ accompanies segregation of free SynGAP and free PSD-95 into complementary regions (Fig. 10).

### 4.1 Model limitations and extensions

The model represents association through a single *S*_3_*P*_2_ complex and represents auxiliary multivalent and higher-order contacts only through effective pairwise Flory–Huggins interaction parameters. It therefore does not resolve binding-site occupancy, network connectivity, or alternative complex stoichiometries. This omission excludes possible coupling between phase separation and percolation or gelation [19].

The model separates equilibrium phase behavior from nonequilibrium spatial dynamics: Flory–Huggins theory identifies phase coexistence, whereas Cahn–Hilliard–reaction dynamics describe the response to a specified parameter perturbation. Phosphorylation, calcium signaling, scaffolding partners, and membrane localization are represented only indirectly by effective parameters.

The reaction closure uses bulk molecular activities and omits square-gradient contributions to the reaction affinity. Consequently, its detailed-balance construction makes the reaction dissipative with respect to the homogeneous bulk free energy but does not guarantee monotonic decay of the full spatial free-energy functional during nonuniform evolution.

We use constant mobility weights to simplify the model and computation; their nondimensionalized values permit the semi-implicit Fourier update to be solved independently for each spatial mode. A composition-dependent choice *m*_*i*_ ∝ *ψ*_*i*_ would provide a degenerate mobility that suppresses transport as species *i* is depleted.

The simulations use idealized periodic domains rather than three-dimensional dendritic-spine geometry. Future work should use reconstructed spine geometries, impose no-flux conditions at the plasma membrane, and explicitly couple the cytosol to the *α* phase surrounding the PSD. Nevertheless, the provisional parameter regime chosen for the Cahn–Hilliard simulations produces spatial and temporal scales compatible with the lateral dimensions of excitatory PSDs and the timescale of acute SynGAP dispersion, including a dissociation timescale of approximately 30–60 s (Fig. 10) [3, 4].

Reduced SynGAP expression alters synaptic structure and function, including dendritic-spine morphology, AMPA-receptor content, and the capacity for subsequent LTP [13]. In the model, Fig. 11 compares equilibrium phase organization at control and reduced Syn-GAP dosage under the pre- and post-LTP parameter sets. For these illustrative parameters, the reduced-SynGAP composition remains within the two-phase region in both conditions, although its tie-line endpoints and phase compositions differ from those of the control composition.

In further work, calibration should compare computed phase compositions and tie-line directions with equilibrium measurements. Time-resolved redistribution data should constrain mobilities and association/dissociation rates. Independent time-resolved experiments could then test the predicted redistribution trajectories, their dependence on SynGAP dosage, and the persistence of a PSD-95-rich phase.

Future extensions could represent the distinct domain structures and binding motifs of N-terminal variants A, B, and C and C-terminal variants *α*_1_, *α*_2_, *β*, and *ϕ*. These isoforms differ in synaptic localization and function: SynGAP-*α*_1_ undergoes phase separation with PSD-95, whereas SynGAP-*β* does not bind the PSD-95 PDZ domains [1]. Earlier work likewise found that SynGAP-*α*_1_- and SynGAP-*α*_2_-containing isoforms exert opposing effects on synaptic strength [18]. Genetic disruption of the endogenous SynGAP-*α*_1_*/α*_2_ sequences in mice also impairs synaptic plasticity and learning and reduces seizure protection [15]. Testing isoform-specific phase behavior, however, would require further investigation of realistic association stoichiometries and interaction parameters.

## 5 Acknowledgments

This work was supported in part by National Science Foundation Grant DMS 1951646 to GDCS. GDCS used the OpenAI Codex desktop application (GPT-5.6 Sol) as an interactive software-development assistant for reproducible figure-generation workflows. All text, mathematics, code, analyses, and figure workflows were conceived by the authors and critically reviewed. The authors take full responsibility for the originality and integrity of the work. Some results appeared previously in the undergraduate thesis of SS [21].

## Appendix A Biological parameterization in physical lattice units

In the Flory–Huggins–Cahn–Hilliard–reaction model, the physical inputs include lattice-site volume, molecular volumes, transport coefficients, interfacial lengths, reaction rates, domain dimensions, and observation times. A nondimensional form is used in the numerical solver, but its dimensionless coefficients are derived from these physical inputs. Lengths and times are reported in nanometers (nm) and seconds (s).

### A.1 Fixed thermal-energy scale

We fix the temperature, *T* = 300 K, for all calculations so that comparisons are not confounded by a change in thermal energy. At this temperature,

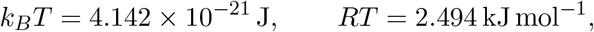

where *R* = *N*_*A*_*k*_*B*_. Thus, an energy written as *αk*_*B*_*T* can be expressed as *α*(4.142 × 10^−21^)J per molecule or *α*(2.494) kJ mol^−1^.

### A.2 Reference lattice-site volume

In the lattice-based Flory–Huggins construction, *v*_0_ denotes the physical volume of one reference lattice site, and *v*_*i*_ denotes the effective molecular volume of species *i*. The corresponding relative-volume parameters are

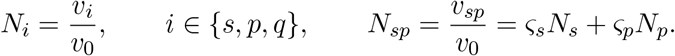

The *N*_*i*_ are dimensionless volume ratios, whereas *v*_0_ and *v*_*i*_ retain physical units. The natural physical bulk-free-energy density scale is

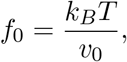

with units J nm^−3^. The physical and nondimensional free-energy densities are related by

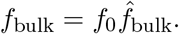

We select the rounded reference-site volume *v*_0_ = 100 nm^3^, motivated by the calculated 97.11 nm^3^ partial volume of one full-length PSD-95 molecule, and define one *Q* unit as one effective site, so that *N*_*p*_ = *N*_*q*_ = 1. This convention does not imply that PSD-95 and water have the same molecular volume; instead, *Q* represents a coarse-grained aqueous-background volume equal to one reference lattice site.

### A.3 Molecular definitions and volume estimates

Here, *P* denotes one PSD-95 molecule, whereas *S* denotes one SynGAP trimer. Accordingly, *v*_*s*_ and *N*_*s*_ refer to the volume and relative volume of an entire SynGAP trimer, not of one SynGAP polypeptide chain. The choices *N*_*s*_ = 1 and *N*_*s*_ = 5 therefore represent a trimer of the SynGAP construct and a full-length SynGAP trimer, respectively, relative to the reference volume assigned to one PSD-95 molecule. We vary the SynGAP-trimer relative volume because the experiments motivating the stoichiometric association model used protein constructs (*N*_*s*_ = 1), whereas the biological interpretation concerns full-length synaptic proteins (*N*_*s*_ = 5).

We estimate molecular partial volumes from polypeptide masses using the standard mean protein partial specific volume 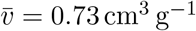 [9],

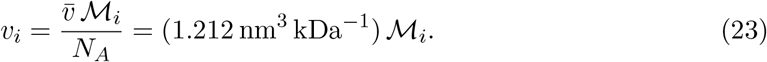

#### In vitro: phase-separating experimental constructs (*N*_*s*_ = *N*_*p*_)

The construct-based species correspond directly to the proteins used by Zeng *et al*. [28] and analyzed by Lin *et al*. [17]:

- *S* is one trimer of the 157-residue SynGAP CC–PBM construct derived from residues A1147–V1308 of mouse SynGAP, with the reported internal deletions. Its calculated sequence mass is 56.575 kDa per trimer, close to the 55.5 kDa solution measurement.
- *P* is one 416-residue human PSD-95 PSG construct spanning R306–L721. Its calculated sequence mass is 47.242 kDa.

Using Eq. 23, the partial-volume estimates are

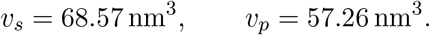

The ratio *v*_*s*_*/v*_*p*_ = 1.20 supports the deliberately low-resolution equal-site approximation *N*_*s*_ = *N*_*p*_ used by Lin *et al*. We therefore adopt the equal-site coarse-graining

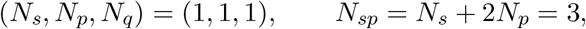

and the fractions of the complex volume attributable to the two protein components are

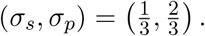

#### In vivo: full-length synaptic proteins (*N*_*s*_ = 5*N*_*p*_)

We define the full-length species as follows:

- *S* is one trimer of the 1308-residue PDZ-binding mouse SynGAP sequence used as the CC–PBM parent (`UniProt J3QQ18`). The calculated mass is 144.866 kDa per monomer and 434.598 kDa per trimer.
- *P* is one full-length 721-residue human PSD-95 molecule from the PSG parent sequence (`RefSeq NP_001122299.1`), with calculated mass 80.125 kDa.

The volume estimates are

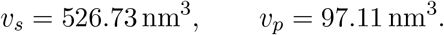

The ratio *v*_*s*_*/v*_*p*_ = 5.42 supports the rounded approximation *N*_*s*_ = 5*N*_*p*_. With *N*_*q*_ = *N*_*p*_, this gives

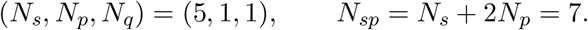

The corresponding complex-volume fractions are

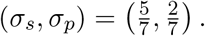

### A.4 Mapping the Lin *et al*. concentrations into the composition simplex

For a homogeneous phase with total component volume fractions *ϕ*_*s*_ and *ϕ*_*p*_, the physical number densities of SynGAP trimers and PSD-95 molecules are *n*_*i*_ = *ϕ*_*i*_*/*(*N*_*i*_*v*_0_), *i* ∈ *{s, p}*, and the corresponding molar concentrations are *c*_*i*_ = *ϕ*_*i*_*/*(*N*_*i*_*v*_0_*N*_*A*_). For *v*_0_ = 100 nm^3^, *c*_*i*_ = (16.61 mM)*ϕ*_*i*_*/N*_*i*_, *i* ∈ *{s, p}*.

Lin *et al*. reported SynGAP concentrations of 30, 60, 90, 120, and 180 *µ*M in monomer-equivalent units [17]. Assuming that the construct resides predominantly in trimers, the corresponding trimer concentrations are *c*^0^ = 10, 20, 30, 40, and 60 *µ*M; the PSD-95 concentrations are 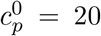, 40, 60, 80, and 120 *µ*M. With the construct-trimer convention *N*_*s*_ = *N*_*p*_ = 1, these SynGAP-trimer and PSD-95 concentration ranges map to

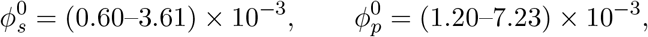

and hence 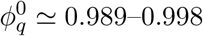. In our ordering ***ϕ*** = (*ϕ*_*s*_, *ϕ*_*p*_, *ϕ*_*q*_), the initial compositions of the experimental mixtures lie close to the solvent-rich vertex (0, 0, 1).

The protein concentrations inside the condensed droplets were much larger than the total concentrations. The reported condensed-phase concentrations correspond to PSD-95 volume fractions 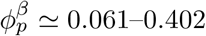 and SynGAP-trimer volume fractions 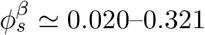.

The low total concentrations and high condensed-phase concentrations are related by the lever rule. If *v*_*β*_ is the fraction of the sample volume occupied by the condensed phase, then

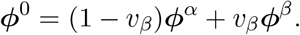

For example, 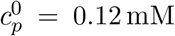 and 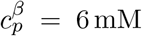 give *v*_*β*_ ≃ 0.02 if the residual dilute-phase concentration is neglected. Thus, reproducing the Lin *et al*. experiments requires a two-phase region that extends close to (0, 0, 1), with a solvent-rich endpoint near the experimental dilute phase and a protein-rich endpoint with volume fractions of order 0.1–0.4.

### A.5 Estimation of species mobility weights

The species mobility weights *m*_*i*_ were estimated from effective collective diffusivities *D*_*i*_, which are more readily related to experimental transport measurements. Because *f*_0_ = *k*_*B*_*T/v*_0_, we use

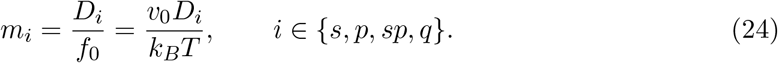

The pairwise mobilities *M*_*i,j*_ for *i ≠ j* and the Onsager matrix *L* then follow from Eqs. 18–19. The *D*_*i*_ have physical dimensions of L^2^*/*T, and the resulting *m*_*i*_ have dimensions of L^5^E^−1^T^−1^.

We anchored the provisional scale at *D*_*p*_ = 0.020 *µ*m^2^ s^−1^ for PSD-95 and estimated the relative protein diffusivities by compact-particle Stokes–Einstein scaling at a common viscosity,

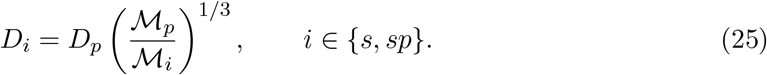

Here the masses are rounded effective values chosen consistently with the relative-volume calibration. For the Fig. 10 calculations, (*N*_*s*_, *N*_*p*_, *N*_*q*_) = (5, 1, 1), so we use (400, 80, 560) kDa. We assign the coarse-grained background the reference value *D*_*q*_ = *D*_*p*_. Eq. 25 then gives

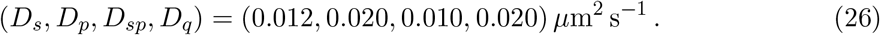

These values are effective collective diffusivities, not measurements of individual SynGAP or PSD-95 molecules. With *v*_0_ = 100 nm^3^, Eq. 24 converts these estimates of *D*_*i*_ to

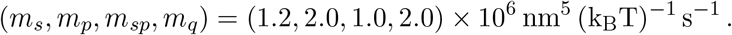

Equivalently, these values are (2.9, 4.8, 2.4, 4.8) × 10^11^*µ*m^5^ J^−1^ s^−1^ for *T* = 300 K.

We note that protein diffusion inside condensates can be substantially slower than suggested by Eq. 26. For example, Taylor *et al*. [23] measured 0.0017 *±* 0.0005 *µ*m^2^ s^−1^ for LAF-1 in its dense phase and showed that diffusivities inferred from fluorescence recovery depend on the transport model and boundary conditions.

## APPENDIX B Concentration- and activity-based kinetic constants

This section explains how concentration-based kinetic and equilibrium parameters map to the activity-based parameters used in the model (Eq. 17). Let *c*_*i*_ denote molar concentration and *v*_*i*_ = *N*_*i*_*v*_0_ the molecular volume assigned to species *i*. These quantities are related to the volume fraction by

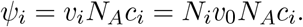

Using *c*_*∘*_ = 1*/*(*v*_0_*N*_*A*_) as the standard concentration gives

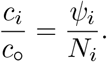

The thermodynamic tendency to participate in a reaction is represented by the dimensionless molecular activity

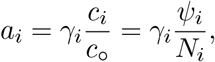

where the dimensionless activity coefficient *γ*_*i*_ accounts for the effect of intermolecular interactions on the thermodynamic availability of species *i*. For an ideal mixture, *γ*_*i*_ = 1, so *a*_*i*_ = *c*_*i*_*/c*_*∘*_ = *ψ*_*i*_*/N*_*i*_. In the present model, *γ*_*i*_ depends on local composition through the Flory–Huggins enthalpic free energy (Eq. 8).

The reversible association in Eq. 4 obeys the concentration-based mass-action equations:

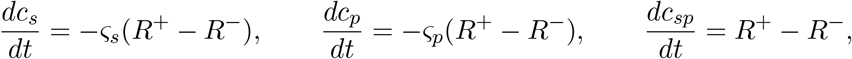

where *R*^+^ and *R*^−^ are the forward and reverse reaction rates per unit volume, respectively:

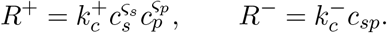

The apparent concentration-based rate coefficients 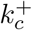 and 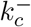 depend on local composition through the activity coefficients. Their physical dimensions are

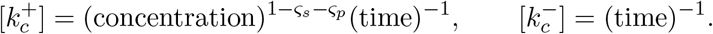

Because *r*_+_ = *R*^+^*/c*_*∘*_ and *r*_−_ = *R*^−^*/c*_*∘*_ have units of inverse time, substituting *a*_*i*_ = *ϕ*_*i*_*c*_*i*_*/c*_*∘*_ into Eq. 17 yields

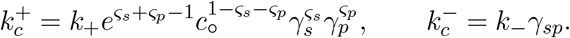

## Appendix C Flory–Huggins interaction energetics

For four species, the Flory–Huggins enthalpy (Eq. 8) contains six dimensionless interaction parameters χ_*ij*_, obtained from ten interaction energies *u*_*ij*_. For simplicity, interactions involving the SynGAP–PSD-95 complex are defined as additive sums of the corresponding constituent interactions. Substitution of Eq. 22 into Eq. 9 gives the map from the six constituent interaction energies (*u*_*ij*_) to six exchange-energy differences (Δ*u*_*ij*_) and six Flory–Huggins interaction parameters (χ_*ij*_ = *z*Δ*u*_*ij*_*/k*_*B*_*T*),

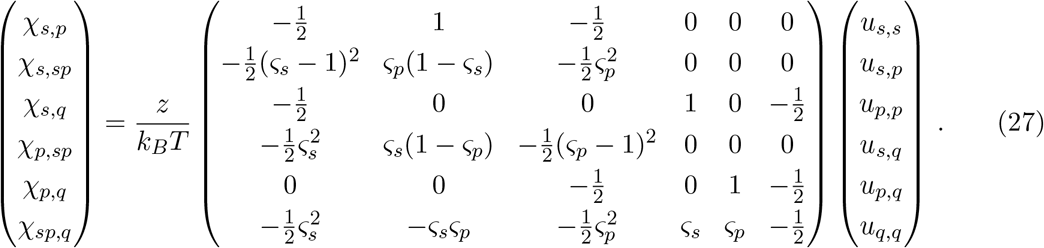

The determinant of this dimensionless coefficient matrix is 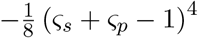. It is therefore nonsingular because both stoichiometric coefficients are integers greater than or equal to 1 and hence *ς*_*s*_ + *ς*_*p*_ *≠* 1. Thus, no nontrivial linear constraint relates the six χ_*ij*_. In fact, for the adopted stoichiometry (*ς*_*s*_, *ς*_*p*_) = (1, 2), Eq. 27 is

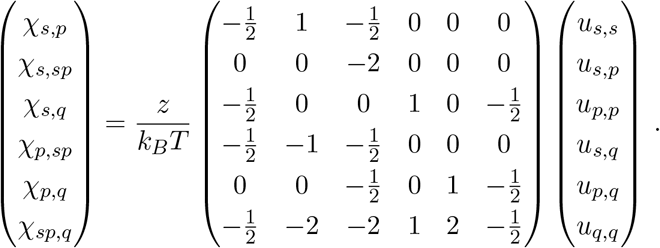

The inverse map is

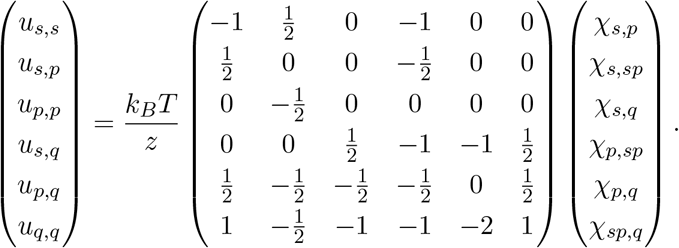

Thus, within the additive-energy subspace for the complex defined by Eq. 22, any six Flory– Huggins parameters (χ_*ij*_) uniquely determine the six constituent interaction energies (*u*_*ij*_).

## Appendix D Physical dimensions and nondimensionalization

This appendix describes the physical dimensions of the Flory–Huggins and Cahn–Hilliard equations and the nondimensionalization used in the simulations. We leave physical quantities unadorned and mark nondimensionalized quantities with hats. Quantities that are dimensionless by definition—including volume fractions, activities, relative molecular volumes, stoichiometric coefficients, and *K*—remain unadorned. We denote the physical dimensions of length, time, and energy by L, T, and E, respectively. These dimensions refer to the underlying three-dimensional continuum; the one- and two-dimensional simulations are spatial reductions of the same scaled equations. For a *d*-dimensional reduction (*d* = 1 or 2), the reduced functional therefore has dimensions EL^*d*−3^, but the functional derivative still has dimensions E*/*L^3^.

### D.1 Dimensional form

Physical position **x** and time *t* have dimensions L and T, respectively, whereas the species volume fractions *ψ*_*i*_ are dimensionless. The three-dimensional free-energy functional

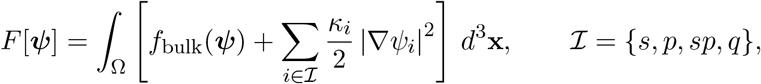

has physical dimensions of energy (E). Because the functional derivative is taken with respect to dimensionless volume fractions, *f*_bulk_ and *µ*_*i*_ = *δF/δψ*_*i*_ both have dimensions of energy density, E*/*L^3^. The dimensional Cahn–Hilliard–reaction equations are

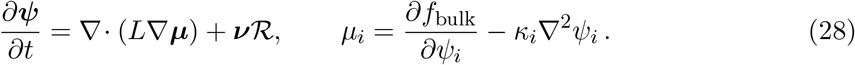

The diffusive fluxes *J*_*i*_ = − ∑_*j*_ *L*_*ij*_*∇µ*_*j*_ have dimensions L*/*T. The pairwise mobilities *M*_*i,j*_ for *i* ≠ *j*, Onsager matrix entries *L*_*ij*_, and species mobility weights *m*_*i*_ all have dimensions L^5^E^−1^T^−1^. Because the activities *a*_*i*_ are dimensionless, the activity-based rates

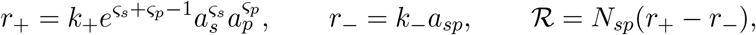

and the rate constants *k*_+_ and *k*_−_ all have dimensions T^−1^.

### D.2 Nondimensionalization

We choose a spatial scale *λ*, a time scale *τ*, and a free-energy-density scale *f*_0_. For the Flory–Huggins lattice model, we use the natural free-energy-density scale

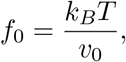

where *v*_0_ is the reference lattice-site volume. We define the dimensionless variables by

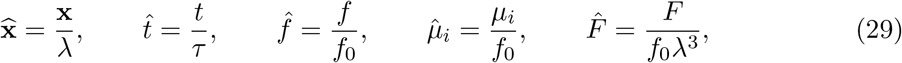

so 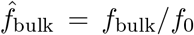 is dimensionless. The dimensionless transport, gradient, and reaction parameters are then

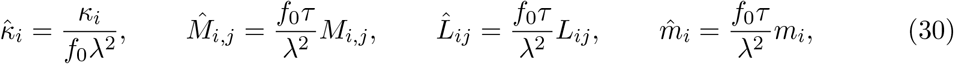

and

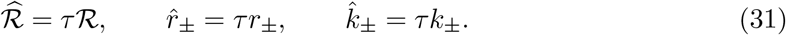

The square-gradient coefficient can equivalently be expressed through the physical gradient length

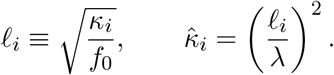

Substituting Eqs. 29–31 into Eq. 28 and using 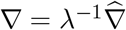 yields

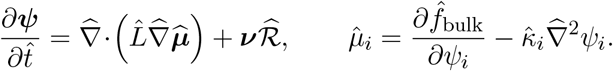

This is the dimensionless system used in the Cahn–Hilliard simulations.

For example, Fig. 10 represents a 1000 nm physical domain by the dimensionless domain length 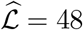. This sets the reference length to

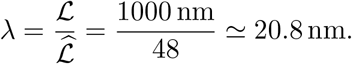

We use the time scale *τ* = 1 s, the reference-site volume *v*_0_ = 100 nm^3^, and hence the free-energy-density scale

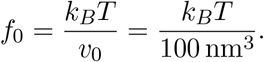

The effective diffusivities estimated in Appendix A.5,

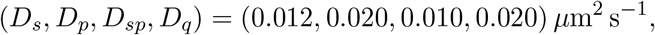

then give

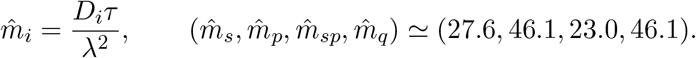

Similarly, the common physical gradient length ℓ_*i*_ = 16 nm gives

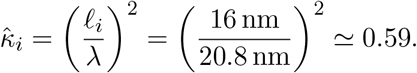

The dimensionless 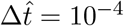 corresponds to the physical time step 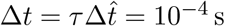.

## Appendix E Semi-implicit Fourier scheme

For clarity, we omit hats from nondimensional quantities throughout this appendix. The simulations use a uniform periodic grid with *N* points in each spatial direction. The nondimensional grid spacing is #*x* = *L/N* . We compute spatial derivatives using discrete Fourier transforms and denote the Fourier coefficient of *φ*(**x**) at wave vector **k** by *F*_**k**_[φ]. With *k* = |**k**|, the Fourier symbols of the Laplacian and biharmonic operators are −*k*^2^ and *k*^4^, respectively. For each coordinate direction, the implementation uses the discrete wave numbers

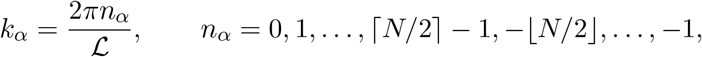

which is the standard periodic FFT ordering for either even or odd *N*.

Let *K* = diag(*κ*_*s*_, *κ*_*p*_, *κ*_*sp*_, *κ*_*q*_). The chemical-potential vector is then

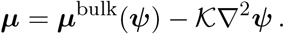

We evaluate the nonlinear bulk chemical potentials and reaction rate explicitly at time step *n* and treat the stiff gradient term implicitly, yielding

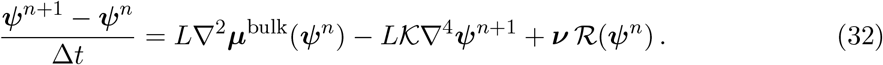

Before solving in Fourier space, we use local incompressibility to eliminate the solvent field. We define 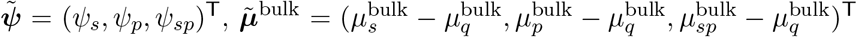, and 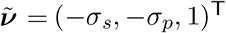. Let 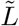 denote the upper-left 3 × 3 block of *L*, as in the main text. Because *L***1** = **0**, the first three rows of the full diffusive update reduce exactly to 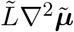. Eliminating *ψ*_*q*_ = 1 − *ψ*_*s*_ − *ψ*_*p*_ − *ψ*_*sp*_ gives the reduced gradient matrix

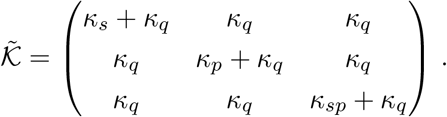

Transforming Eq. 32 to Fourier space yields the following 3 × 3 linear system for each mode:

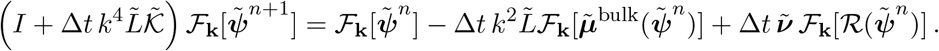

Each Fourier mode requires the solution of a 3 × 3 linear system. The implementation constructs and caches its inverse for each distinct trial substep size and applies the cached inverse at each accepted substep, after which local incompressibility determines *ψ*_*q*_. For the zero-wave-number mode, diffusion vanishes and the spatially averaged reaction changes the individual species means according to

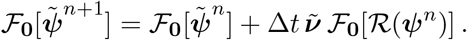

We define ***ν***_*s*_ = (1, 0, *σ*_*s*_, 0)^⊤^ and ***ν***_*p*_ = (0, 1, *σ*_*p*_, 0)^⊤^. Then 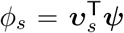 and 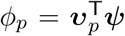. The identities 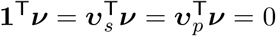 ensure conservation of total volume, SynGAP, and PSD-95 despite free–bound interconversion.

Here, Δ*t* is the current nondimensional trial substep for the four-species calculation. The algorithm begins each nominal time interval with the configured step size. If a candidate contains a nonfinite species fraction or lies outside the four-species physical simplex, the algorithm discards the trial, halves the step size, and retries from the last accepted state. Accepted dyadic substeps partition the nominal interval. The scheme is first-order accurate in time and is not unconditionally energy stable.

For the no-association calculations in Figs. 2 and 3, the reaction term vanishes. We set *m*_*s*_ = *m*_*p*_ = *m*_*q*_ = 1, giving *M*_*i,j*_ = 1*/*3 for distinct *i, j* ∈ *{s, p, q}*, and 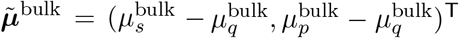. The resulting scheme evolves the two fields 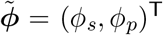 and treats the full coupled gradient block implicitly:

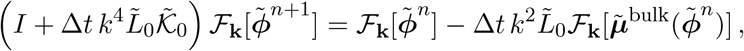

where

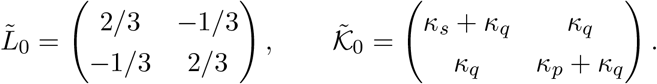

The zero-wave-number coefficients are copied unchanged, so the spatial means of *ϕ*_*s*_ and *ϕ*_*p*_ are preserved to roundoff.

## Appendix F Identifying binodals, spinodals, and association-branch switch loci

At fixed total composition ***ϕ*** = (*ϕ*_*s*_, *ϕ*_*p*_, *ϕ*_*q*_), we define 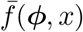 as the unminimized nondimensional free-energy density with association coordinate *x* = *ψ*_*sp*_. Writing 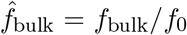, its definition in the independent coordinates (*ϕ*_*s*_, *ϕ*_*p*_) is

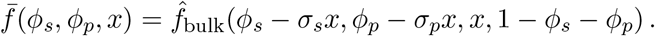

Here, *f*_bulk_(*ψ*_*s*_, *ψ*_*p*_, *ψ*_*sp*_, *ψ*_*q*_) is given by Eqs. 6–10. A nondegenerate local minimum *x*_*ψ*_(***ϕ***) satisfies 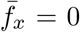 and 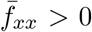. It defines a locally stable association branch with free energy 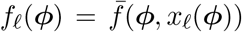. The equilibrium reduced free energy is the pointwise minimum over these branches:

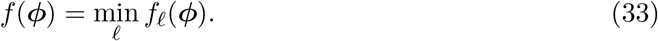

Consequently, a branch can be locally stable along the association coordinate but metastable relative to another branch. The reduced free energy is smooth within each selected branch but can be nondifferentiable where the global minimum switches branches. The domain is the composition simplex

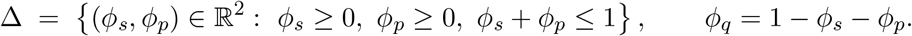

Throughout this appendix, ***ϕ*** ∈ Δ, whether used as an argument of *f* or in a dot product with ***µ***, denotes the independent pair (*ϕ*_*s*_, *ϕ*_*p*_), with *ϕ*_*q*_ = 1 − *ϕ*_*s*_ − *ϕ*_*p*_. We determine phase-coexistence regions and binodal boundaries by lower-convex-hull construction (§F.2) and use constrained sampling of lever-rule-compatible two-phase states at selected compositions as an independent numerical check (§F.1).

### F.1 Algebraic construction of the two-phase free energy

For a prescribed overall composition ***ϕ***_0_, the phase compositions and volume fractions satisfy the lever rule

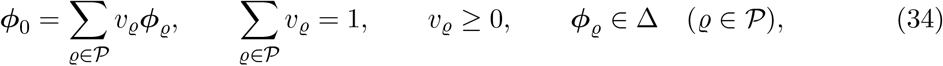

Here, *P* = *{α, β}* for two-phase coexistence and *P* = *{α, β, γ }* for three-phase coexistence. In Eq. 34 and throughout this appendix, subscripts label phases.

For a two-phase decomposition with compositions ***ϕ***_*α*_ and ***ϕ***_*β*_, the mixture free-energy density is

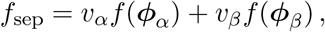

with *v*_*α*_ + *v*_*β*_ = 1 [20]. Eliminating *v*_*α*_ and ***ϕ***_ε_ using Eq. 34 yields

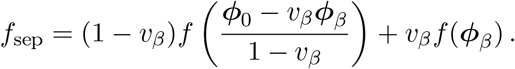

The implementation samples a candidate phase composition and volume fraction, either randomly or on a grid, and reconstructs the other phase by the lever rule. The smallest sampled value of *f*_sep_ estimates the lowest two-phase free-energy density at ***ϕ***_0_, and its associated phase pair approximates a supporting line segment of the convex envelope. This constrained-sampling calculation is an independent numerical check of the lower-hull construction.

### F.2 Convex-hull characterization

Convexification of *f*(***ϕ***) over Δ determines equilibrium phase separation. The coarse-grained equilibrium free energy is the convex envelope

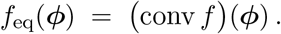

A composition lies in the phase-coexistence region when

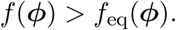

Numerically, the lower convex hull of the sampled surface *{*(***ϕ***,*f*(***ϕ***))*}* is represented by planar supporting facets. We use MATLAB’s `convhulln` to compute the convex hull of the sampled points (***ϕ***,*f*(***ϕ***)). We retain facets whose outward normals have a negative free-energy component. These lower-hull facets discretize the convex envelope.

#### Binodal: common-supporting-plane conditions

Two compositions ***ϕ***_ε_ and ***ϕ***_*β*_ coexist when they share a supporting plane

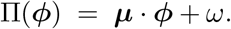

This plane touches *f* at both compositions and nowhere exceeds *f* on Δ:

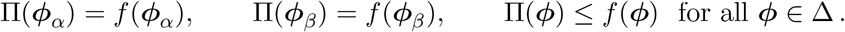

We use the independent coordinates (*ϕ*_*s*_, *ϕ*_*p*_), with *ϕ*_*q*_ = 1 − *ϕ*_*s*_ − *ϕ*_*p*_. At differentiable contact points, the binodal conditions are

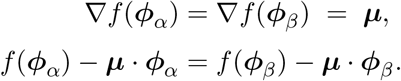

Here ***µ*** is the vector of solvent-referenced reduced chemical potentials associated with (*ϕ*_*s*_, *ϕ*_*p*_). It determines the slope of the supporting plane. The common intercept 5 = *f* − ***µ*** · ***ϕ*** is the reduced semigrand-potential density. The contact compositions lie on the binodal boundaries of the ternary composition diagram, and the segments joining coexisting pairs are tie lines.

Three-phase coexistence occurs when one supporting plane touches *f* at three distinct compositions ***ϕ***_ε_, ***ϕ***_*β*_, and ***ϕ***_*γ*_. If *f* is differentiable at these contact points, the coexistence conditions are

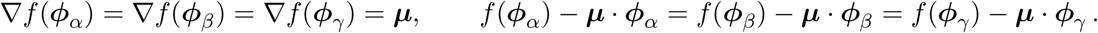

These equalities define a flat triangular facet of the convex envelope (Fig. 1B).

#### Spinodal: loss of local convexity

Unlike the binodal, the spinodal is a local stability boundary where the Hessian of *f*, restricted to Δ, loses positive definiteness. On a selected smooth association branch, we set *x* = *ψ*_*sp*_ and partition the 3 × 3 Hessian of 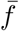 between the conserved coordinates ***ϕ*** = (*ϕ*_*s*_, *ϕ*_*p*_) and *x*:

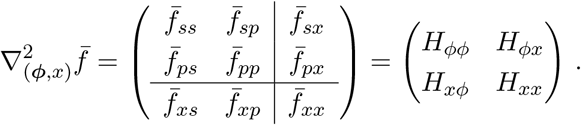

Subscripts on 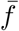 denote differentiation with respect to (*ϕ*_*s*_, *ϕ*_*p*_, *x*). For example, 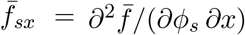. Here, *H*_*ϕϕ*_ is the 2 × 2 composition block, *H*_*xx*_ is the scalar association block, and *H*_*ϕx*_ and *H*_*xϕ*_ are the mixed blocks. When *H*_*xx*_ > 0, the stationary root is a nondegenerate local minimum in *x*. Minimizing over *x* then gives the reduced Hessian as the Schur complement

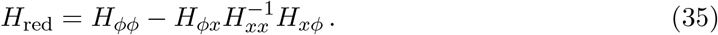

We define the dimensionless species-coordinate Hessian entries by

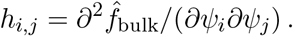

For the Flory–Huggins free energy in Eqs. 7–8,

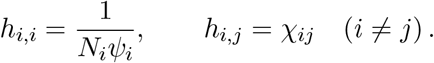

Because the reaction-energy term is linear in *x*, it contributes no second derivatives. The six independent second derivatives of 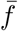 needed in Eq. 35 are

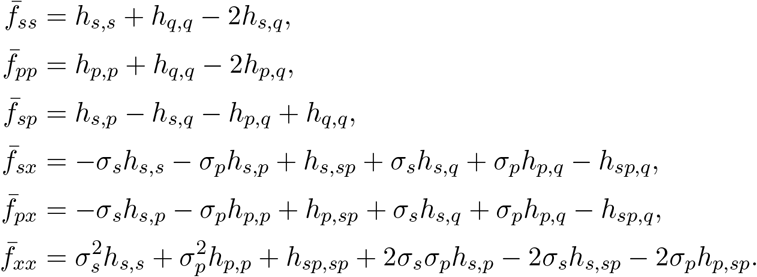

Subscripts on 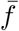 refer to (*s, p, x*) = (*ϕ*_*s*_, *ϕ*_*p*_, *x*), whereas subscripts on *h* refer to the species coordinates (*s, p, sp, q*) = (*ψ*_*s*_, *ψ*_*p*_, *ψ*_*sp*_, *ψ*_*q*_). Expanding Eq. 35 gives

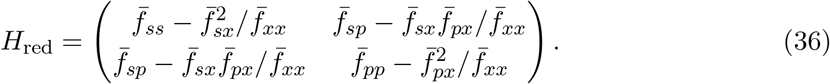

For each phase diagram, we evaluate the analytic Hessian at every composition-grid point and identify the spinodal by the contour *λ*_min_ = 0. Eq. 36 is always evaluated on the globally minimizing association branch (Eq. 33). At a branch-switch locus, the reduced free energy is nondifferentiable; the smooth-branch spinodal therefore terminates there, and we plot the switch locus separately. We verified the analytic Hessian independently against centered finite differences in a smooth test regime confined to a single association branch.

## Appendix G Multiple association stationary roots

With ideal activities, the mass-action equation has one admissible root: an interior root when *ϕ*_*s*_, *ϕ*_*p*_ > 0 and the boundary state *x* = 0 otherwise. Nonideal Flory–Huggins activities can render the constrained free energy nonconvex and produce multiple roots. We enumerate these roots, classify them by the constrained free-energy curvature, and select the global minimizer as the equilibrium complex fraction.

At fixed *ϕ*_*s*_ and *ϕ*_*p*_, let *x* = *ψ*_*sp*_. The remaining species fractions and admissible interval are

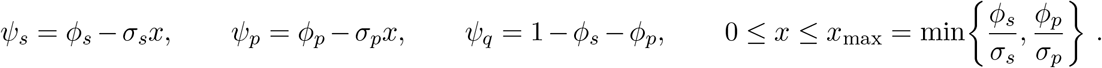

With *f*_0_ = *k*_*B*_*T/v*_0_, define 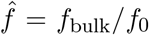. The constrained dimensionless free-energy density is

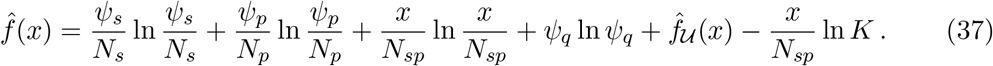

Stationary roots satisfy Λ (*x*) = 0, where 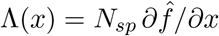:

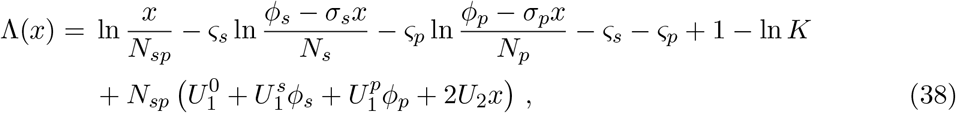

where we define the interaction combinations

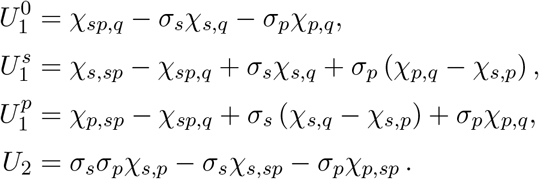

Differentiation with respect to *x* gives

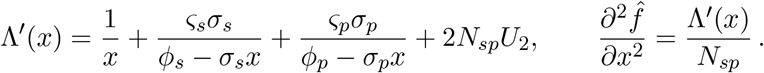

Because *N*_*sp*_ > 0, Λ′(*x*) and the constrained free-energy curvature have the same sign; thus, Λ′ (*x*) classifies the stationary roots. When *U*_2_ = 0, every term in Λ′ (*x*) is positive. For *K* > 0 and *ϕ*_*s*_, *ϕ*_*p*_ > 0, ′ (*x*) increases from − ∞ as *x ⟶* 0^+^ to +∞ as 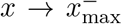. It is therefore strictly increasing and has exactly one admissible interior root.

Nonideal interactions permit three stationary roots only when *U*_2_ is sufficiently negative to make Λ′ (*x*) < 0 over an intermediate interval, producing an S-shaped mass-action residual. If the local maximum is positive and the local minimum is negative, & crosses zero three times: the two outer roots are local free-energy minima, and the middle root is a local maximum.

As an illustration, we use *K* = 10^3^, *u*_*ss*_ = 1*/*10, *u*_*sp*_ = −3*/*20, and *u*_*pp*_ = −1*/*20, with *k*_*B*_*T* = 1 and *z* = 6. We set the solvent interaction energies to zero. The molecular volumes are (*N*_*s*_, *N*_*p*_) = (5, 1), and the stoichiometric coefficients are (*ς*_*s*_, *ς*_*p*_) = (1, 2). These choices give *N*_*sp*_ = 7, (*σ*_*s*_, *σ*_*p*_) = (5*/*7, 2*/*7), and

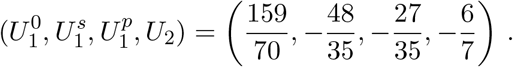

On the 180-subdivision simplex grid, the fold loci are traced as zero contours of &(*x*) at the two roots of Λ′(*x*) = 0 and bound the three-root region (Fig. 12A). At the marked composition (*ϕ*_*s*_, *ϕ*_*p*_, *ϕ*_*q*_) = (0.4, 0.4, 0.2), *x*_max_ = 0.56. Panel B shows the mass-action residual and constrained free-energy landscape at this composition.

**Figure 12.**
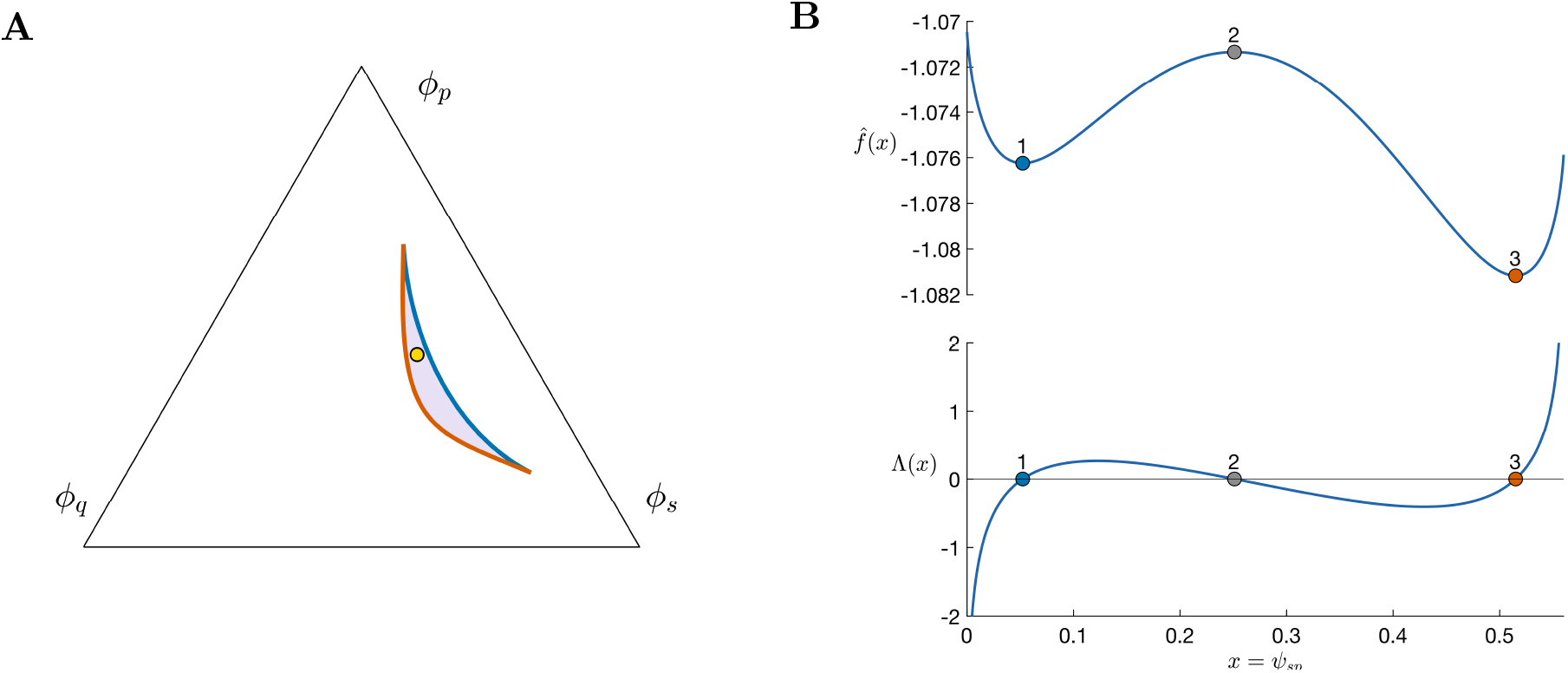
Association stationary-root landscape. **A**: On the 180-subdivision simplex grid, saddle-node loci are zero contours of Λ(*x*) at the roots of Λ′ (*x*) = 0 (*x* = *ψ*_*sp*_). Along the blue fold, the low-*x* minimum coalesces with the intervening maximum; along the orange fold, the maximum coalesces with the high-*x* minimum. In the interior, the region between the folds has three roots and the region outside has one; reactant-free edges have *x* = 0, and each fold has one coalesced pair. The yellow circle marks (*ϕ*_*s*_, *ϕ*_*p*_, *ϕ*_*q*_) = (0.4, 0.4, 0.2). **B**: Constrained free energy 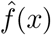 (upper, Eq. 37) and mass-action residual Λ(*x*) (lower, Eq. 38) at the composition marked in panel **A.** The three numbered stationary roots comprise two local minima separated by a local maximum; root 3 is the global minimum.

